# The RNA of maize chlorotic mottle virus - the essential virus in maize lethal necrosis disease - is translated via a panicum mosaic virus-like cap-independent translation element

**DOI:** 10.1101/2020.01.08.898825

**Authors:** Elizabeth Carino, Kay Scheets, W. Allen Miller

## Abstract

Maize chlorotic mottle virus (MCMV) combines with a potyvirus in maize lethal necrosis disease (MLND), an emerging disease worldwide that often causes catastrophic yield loss. To inform resistance strategies, we characterized the translation initiation mechanism of MCMV. We report that, like other tombusvirids, MCMV RNA contains a cap-independent translation element (CITE) in its 3’ untranslated region (UTR). The MCMV 3’ CITE (MTE) was mapped to nucleotides 4164-4333 in the genomic RNA. SHAPE probing revealed that the MTE is a variant of the panicum mosaic virus-like 3’ CITE (PTE). Like the PTE, electrophoretic mobility shift assays (EMSAs) indicated that eukaryotic translation initiation factor 4E (eIF4E) binds the MTE despite the absence of a m^7^GpppN cap structure, which is normally required for eIF4E to bind RNA. The MTE interaction with eIF4E suggests eIF4E may be a soft target for engineered resistance to MCMV. Using a luciferase reporter system, mutagenesis to disrupt and restore base pairing revealed that the MTE interacts with the 5’ UTRs of both genomic RNA and the 3’-coterminal subgenomic RNA1 via long-distance kissing stem-loop base pairing to facilitate translation in wheat germ extract and in protoplasts. However, the MTE is a relatively weak stimulator of translation and has a weak, if any, pseudoknot, which is present in the most active PTEs. Most mutations designed to form a pseudoknot decreased translation activity. Mutations in the viral genome that reduced or restored translation prevented and restored virus replication, respectively, in maize protoplasts and in plants. We propose that MCMV, and some other positive strand RNA viruses, favors a weak translation element to allow highly efficient viral RNA synthesis.

**Author Summary:** In recent years, maize lethal necrosis disease has caused massive crop losses in East Africa and Ecuador. It has also emerged in East Asia. Maize chlorotic mottle virus (MCMV) infection is required for this disease. While some tolerant maize lines have been identified, there are no known resistance genes that confer full immunity to MCMV. In order to design better resistance strategies against MCMV, we focused on how the MCMV genome is translated, the first step of gene expression required for infection by all positive strand RNA viruses. We identified a structure (cap-independent translation element) in the 3’ untranslated region of the viral RNA genome that allows the virus to usurp a host translation initiation factor in a way that differs from host mRNA interactions with the translational machinery. This difference may guide engineering of – or breeding for – resistance to MCMV. Moreover, this work adds to the diversity of known eukaryotic translation initiation mechanisms, as it provides more information on mRNA structural features that permit noncanonical interaction with a translation factor. Finally, owing to the conflict between ribosomes translating and viral replicase copying viral RNA, we propose that MCMV has evolved a relatively weak translation element in order to permit highly efficient RNA synthesis, and that this replication-translation trade-off may apply to other positive strand RNA viruses.

## Introduction

Maize lethal necrosis disease (MLND, also referred to as corn lethal necrosis) first identified in the Americas in the 1970’s [1], has recently spread worldwide, causing devastating crop losses and food insecurity across East Africa, where maize is the most important subsistence and cash crop [2–9]. It has also emerged in China [10], Taiwan [11], Spain [12], and Ecuador, where the damage was so catastrophic in 2015 and 2016 that a state of emergency was declared [13, 14]

MLND is caused by a mixed infection of maize chlorotic mottle virus (MCMV) and any potyvirus that infects maize, usually sugarcane mosaic virus (SCMV) [1, 15, 16]. However, MCMV infection can also be severe when combined with abiotic stress such as drought, or with viruses outside the Potyviridae family [9], while common maize potyviruses like SCMV are generally mild on their own [15, 17, 18]. Efforts to identify genetic resistance against MCMV and potyviruses have revealed resistance to SCMV [19–22], but only tolerance to MCMV [23] with reduced symptoms and virus levels. To our knowledge, no genes that confer complete resistance to MCMV have been identified. Despite its economic importance [5, 24–26], little is known about the molecular mechanisms of MCMV replication, gene expression or its interactions with the host, which could provide valuable knowledge toward identifying targets for resistance breeding or engineering strategies.

MCMV is the sole member of genus *Machlomovirus* in the family *Tombusviridae* [27]. The 4437-nucleotide (nt) positive sense RNA genome contains no 5’ cap, no poly(A) tail, and encodes seven open reading frames (ORFs) [28–30]. The 5’ end of the genome contains two overlapping ORFs that code for a 32 kDa protein (P32) and a 50 kDa (P50) replicase-associated protein (RAP). The P50 ORF has a leaky stop codon which allows for readthrough translation of a 61 kDa C-terminal extension on P50 to form the 111 kDa RNA-dependent RNA polymerase (RdRp) [31] (Fig 1A). In infected cells, MCMV generates two 3’-coterminal subgenomic RNAs that are 5’-truncated versions of the genomic RNA. Subgenomic RNA1 (sgRNA1), spanning nts 2971-4437 serves as mRNA from which the coat protein (CP), and the movement proteins P7a, P7b and P31 are translated. The 337 nt sgRNA2, representing the 3’ untranslated region (UTR), is a noncoding RNA [28]. Although the MCMV genome has been characterized to some extent [17, 28, 31], little is known about its translation mechanisms, a key process in the replication cycle.

**Fig 1.**
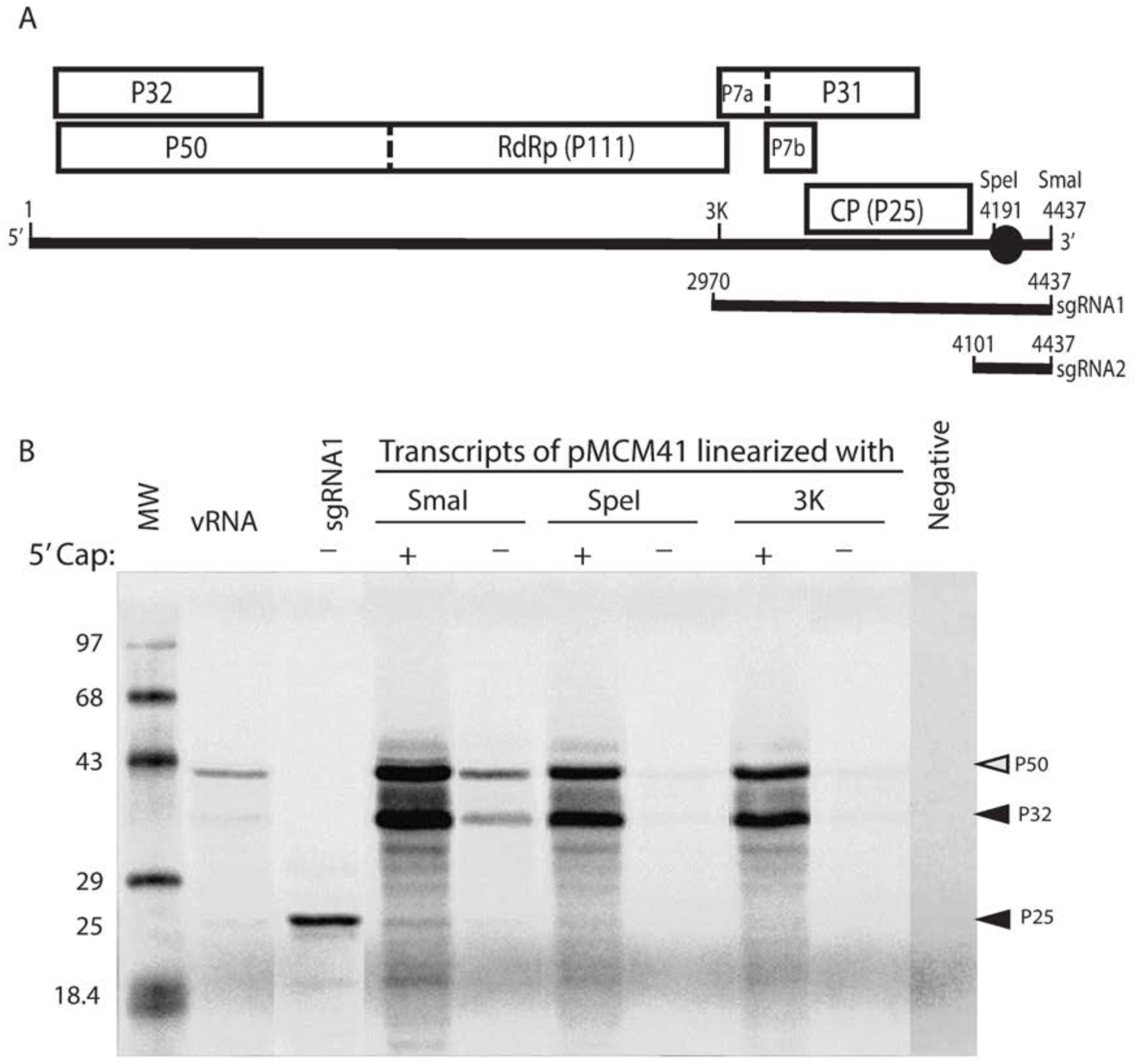
*In vitro* translation of MCMV genomic RNA. (A) Genome organization of MCMV. Boxes indicate open reading frames (ORFs) with encoded protein (named by its molecular weight in kDa) indicated. Dashed boundaries indicate leaky stop codons. RdRp: RNA-dependent RNA polymerase domain of P111; CP: coat protein. Positions of key restriction enzymes are indicated (Sma I: 4437, Spe I: 4191). Black oval indicates location of the 3’ CITE (nts 4164-4333). (B) Translation of capped (+) and uncapped (−) transcripts and virion RNA. pMCM41 was linearized at the indicated locations in the genome prior to transcription. Capped transcripts were from pMCM721, which is identical to pMCM41 but with one nonviral G at the 5’ end, which was required for capping. sgRNA1 is uncapped transcript from psgRNA1 linearized with Sma I. Uncapped transcript from Sma I-linearized pMCM41 is infectious [29]. sgRNA1 is an uncapped in vitro transcript from p Gel shows ^35^S-met-labeled products translated in WGE. The prominent bands identified on the gel correspond to P32 and P50 products (See ORFs in Fig. 1A).

Many positive strand RNA viruses use noncanonical translation mechanisms, including cap-independent translation. This frees the virus from having to encode capping enzymes, and also allows the virus to avoid the host’s translational control system which often acts through cap-binding proteins [32–34]. Because it differs from host mechanisms, this virus-specific translation mechanism may provide unique targets for antiviral strategies. A translation strategy used by all studied tombusvirids is to harbor a cap-independent translation element (CITE) in the 3’ UTR of the virus’ genomic RNA, which is uncapped [35–37]. The 3’ CITE replaces the role of the m^7^GpppN cap structure present at the 5’ end of all eukaryotic mRNAs. About seven different structural classes of 3’ CITE are known [35, 38, 39]. Most 3’ CITEs attract the key ribosome-recruiting eukaryotic translation initiation factor heterodimer, eIF4F, by binding to one or both of its subunits, eIF4G or eIF4E [35, 39–43].

Because all other tested tombusvirids harbor a 3’ CITE, we predicted that MCMV RNA harbors a 3’ CITE to facilitate its translation. *In silico* analysis of the MCMV 3’ UTR using MFOLD and ViennaRNAfold to predict RNA secondary structures did not reveal a structure resembling a known 3’ CITE. Here we provide experimental data that demonstrate the presence and function of a 3’ CITE, that we call the MCMV 3’ CITE (MTE). The MTE is structurally similar to the panicum-mosaic virus-like translation element (PTE) class of CITE. We identify a key translation initiation factor with which the MTE interacts (eIF4E), and show how the MTE base pairs to the 5’ UTR to facilitate cap-independent translation, and that the functional MTE and the long-distance interaction are required for infection of maize. The results contribute to our understanding of structure-function relationships of cap-independent translation elements, and valuable information on the first step of gene expression of an important pathogen.

## Results

### Mapping the 3’-cap independent translation element in MCMV

To roughly map the 3’ CITE of MCMV, we translated 3’-truncated transcripts from a full-length cDNA clone of the MCMV genome (pMCM41 [29]). pMCM41 DNA templates were transcribed in the absence of cap analog while pMCM721, which lacks the 5’ terminal A of the MCMV genome, was used for capped transcripts, because the 5’ A of pMCM41 (identical to MCMV RNA) cannot be capped using an m^7^GpppA cap analog and T7 RNA polymerase (KS unpublished observation). Plasmid psgRNA1 was the template for transcription of full-length sgRNA1, the mRNA for the 25 kDa CP and the movement proteins [28]. Transcribed RNAs and RNA isolated from virions (vRNA) were translated in wheat germ extract (WGE) in the presence of ^35^S-methionine. The full-length, infectious transcript from SmaI-linearized pMCM41 and vRNA yielded two protein products, P32 and P50 from the 5’-proximal overlapping ORFs (Fig 1B). Interestingly, vRNA yielded much less P32 protein, relative to P50, than did the transcribed mRNA. Also, a faint band comigrating with CP is visible from both vRNA and the full-length transcript. The expected 111 kDa protein generated by readthrough of P50 stop codon was not detected, most likely because ribosomal readthrough occurs at a very low rate in these conditions. Readthrough products have been difficult to detect among the in vitro translation products of other tombusvirid genomes as well [44–46]. Unlike the full-length genomic RNA from SmaI-cut pMCM41, which yielded substantial protein products, the uncapped 3’-truncated transcripts produced almost no detectable protein product, suggesting that the 3’ CITE is downstream of the Spe I site at nt 4191 (Fig 1B). It is noteworthy that translation in the presence of a 5’ cap on full-length and truncated pMCM41-derived RNAs gave much more translation product than uncapped full-length pMCM41 transcript or vRNA, indicating that the viral genome may be a relatively inefficient mRNA.

To rapidly map the 3’ CITE location at high resolution, a luciferase reporter (MlucM) was constructed such that the coding region of the virus was replaced by the firefly luciferase (Fluc) coding sequence (Fig 2A). Deletion analyses showed little decrease in translation *in vitro* or *in vivo* when either the 3’-terminal 104 nt (nts 4334-4437) or the first 169 nt (nts 4095-4263) of the 3’ UTR were deleted (Fig 2B). Additional constructs that included the adjacent sequence upstream of the 3’ UTR, up to nt 3578 in the CP ORF, translated more efficiently than those that contained only the 3’ UTR (S1 Fig). However, the sequence upstream of the 3’ UTR (3578-4108) alone was not enough to support translation, and the greatest contributor to translation was mapped to the 3’ UTR. Numerous deletions in the MCMV 3’ UTR revealed that the region between nucleotides 4164-4333 produced luciferase activity >100% of that from the full-length 3’ UTR *in vitro* and about 50% *in vivo*. The lower level of translation *in vivo* may be due to reduced RNA stability owing to the absence of the 3’ end, which is thought to confer stability in related viruses because of its highly base-paired terminal bases [47–49]. Deletions within nts 4164 to 4333, especially of nts 4200-4300, reduced luciferase translation *in vitro,* so in subsequent studies we focused on nts 4164-4333 to characterize the MCMV 3’ CITE (MTE).

**Fig 2.**
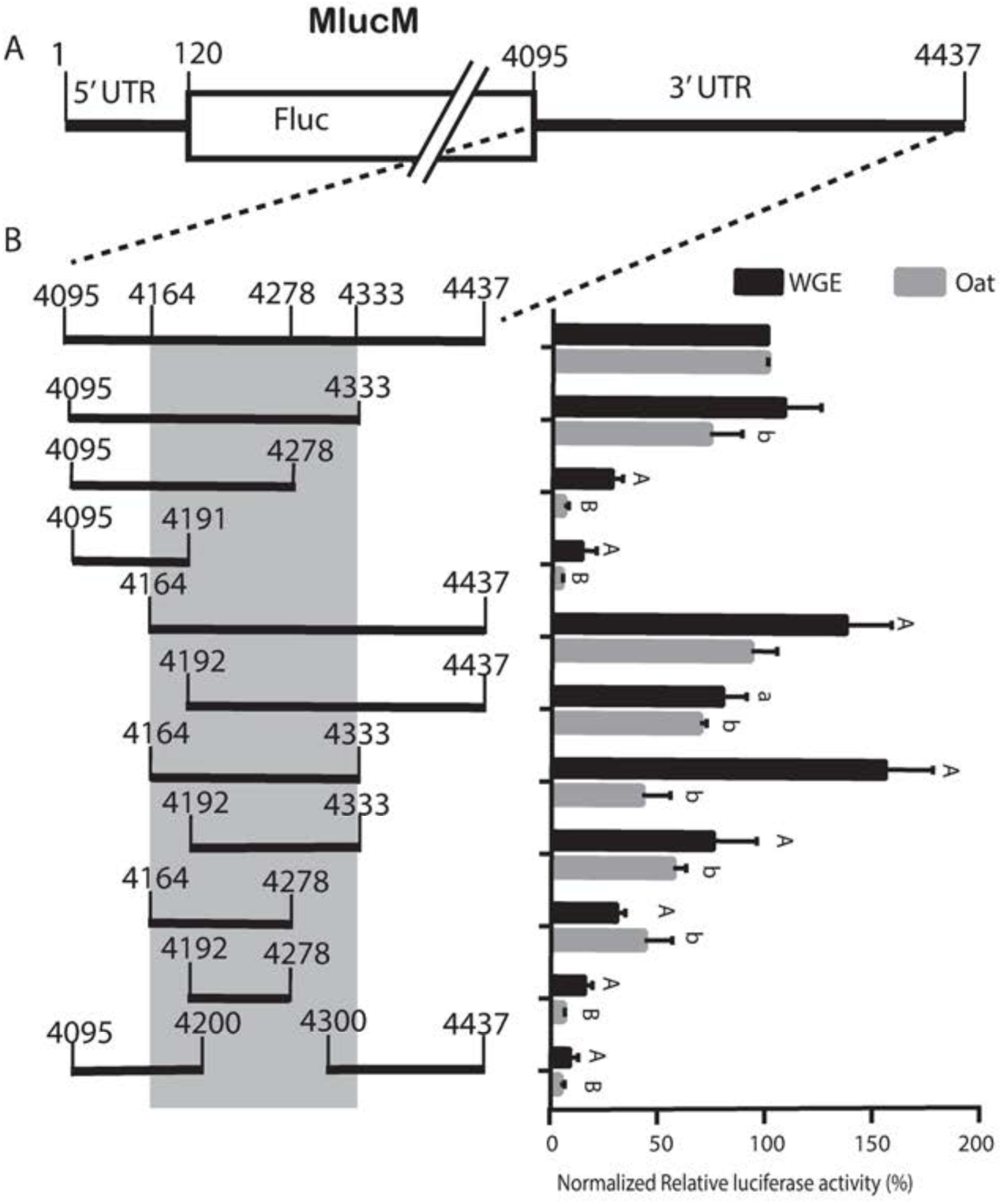
Mapping the sequence in MCMV 3’UTR required for cap-independent translation. (A) Map of MlucM containing a firefly luciferase reporter gene (Fluc) flanked by the complete MCMV UTRs, with base numbering as in the full-length genome. (B) Black bars below left indicate the portions of the 3’ UTR present in each MlucM deletion construct. Sequence covering nt 4164-4333 (grey shading) shows the minimal amount of sequence required for efficient cap-independent translation. Relative luciferase activity obtained from indicated uncapped transcripts in WGE and oat protoplasts is shown as percentages of relative light units where constructs were normalized to MlucM wild type 3’ UTR (top bar). Data are shown as averages, relative to full-length 3’ UTR (±S.D.), from 4 independent experiments (for each construct: WGE: n=10, Oats: n=14). Protoplasts were co-electroporated with the indicated uncapped transcript plus capped Renilla luciferase mRNA as an electroporation control. Fluc/Rluc sample ratios were normalized to MlucM_wt_. One-way ANOVA was used to analyze the significance of each set of samples. For Dunnett’s multiple comparison test significance, an “uppercase” was assigned to a construct if the difference between MlucMwt and deletion mutant was P<0.001 whereas a “lowercase” was given to P<0.05. “A or a” was assigned to data comparisons against MlucMwt in WGE while “B or b” was assigned to comparisons in oat protoplasts.

### Determining the secondary structure of MCMV 3’-CITE

To determine the structure of the MTE, we first attempted to predict its secondary structure computationally. Two different algorithms, MFOLD [50], and ViennaRNA Package [51], predicted a hammer-shaped structure unlike any known 3’ CITE, so we proceeded to determine its secondary structure experimentally by subjecting the MTE (nts 4164-4333) to selective 2’-hydroxyl acylation analyzed by primer extension (SHAPE) probing (Fig 3A). This revealed that the MTE consists of a long helix with various asymmetric internal loops topped by two branching stem loops (Fig 3B), which differed from the computer-predicted structure. The main stem contains a purine-rich bulge between nucleotides 4216-4223. In the presence of magnesium ion, bases G_4215,_ A_4216_, and G_4219_ were hypermodified by the SHAPE reagent benzoyl cyanide, while bases AGA_4221-4223_ became hypomodified (Fig 3A). The MTE also contains a single-stranded “bridging domain” (nts 4246-4250) connecting the two branching stem-loops, which was moderately modified in the presence and absence of magnesium. Side loop-I (SL-I_4235:4241_) houses a pentamer, UGCCA_4236-4240_, in its loop that is complementary to sequence UGGCA in the 5’-UTR. These pentamers may create a long-distance base-pairing interaction between the 3’ and 5’ UTRs (discussed later). The overall structure obtained from the SHAPE probing assays indicates that the MTE has a similar structure to those of Panicum mosaic virus-like 3’ CITES (PTEs) [38, 52], but differing by the presence of three hypermodified bases rather than a single hypermodified G in the purine-rich bulge in the presence of Mg^2+^ [53].

**Fig 3.**
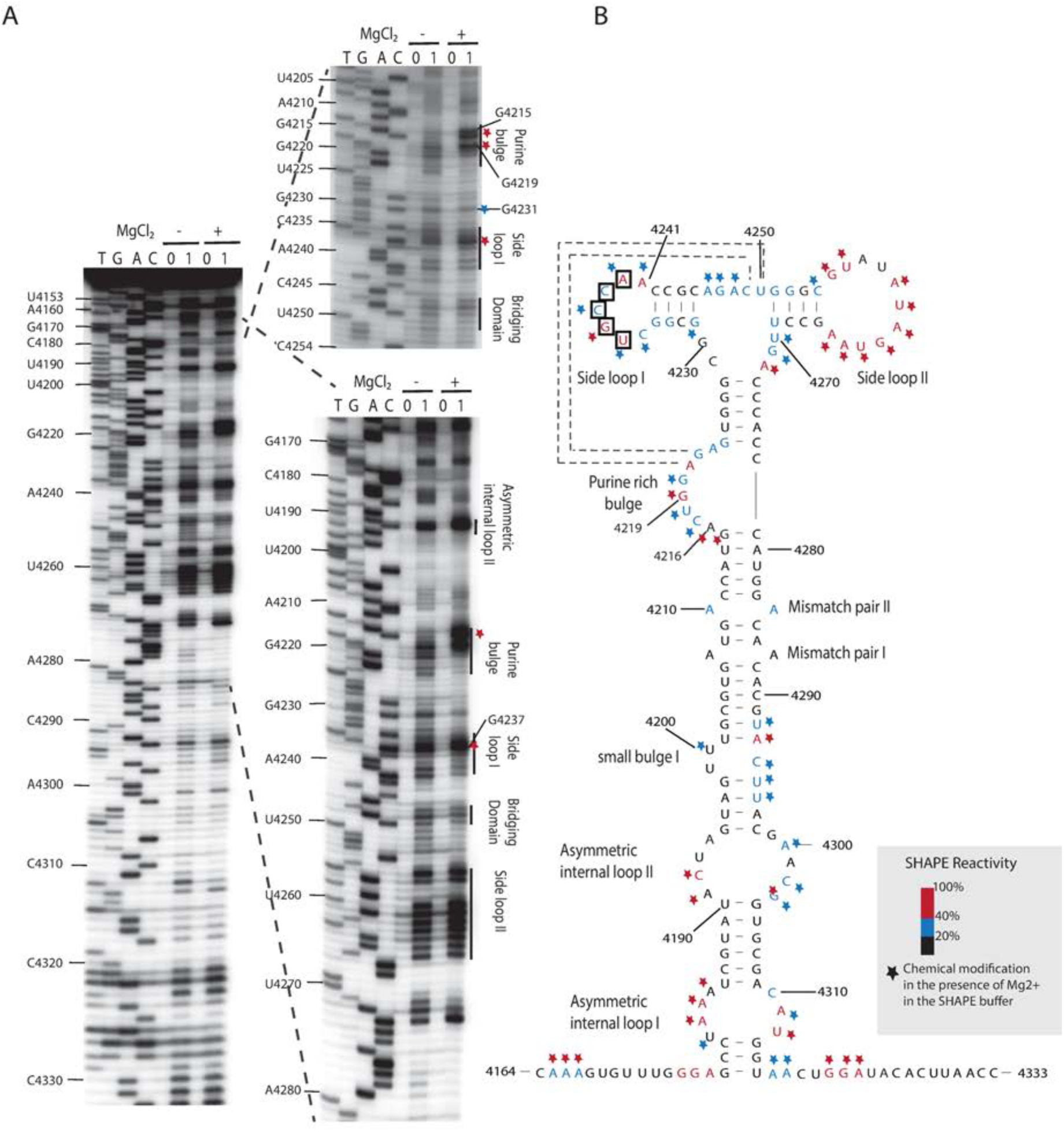
SHAPE analysis of MTE structure from the MCM41 infectious clone. (A) SHAPE (selective 2’-hydroxyl acylation analyzed by primer extension) probing gel. RNA was modified with either 60 mM benzoyl cyanide (1) in dimethyl sulfoxide (DMSO) or DMSO only (0) in the presence (+) or absence (−) of magnesium in SHAPE buffer. (See Methods for details.) Gels show the products of reverse transcriptase extension of ^32^P-labeled primer on SHAPE-modified (1) and unmodified (0) templates. The sequencing ladders (four lanes at left, TGAC) were generated by dideoxy sequencing of unmodified RNA with the same 5’-labeled primer used in the modification lanes. Mobilities of selected nucleotides, numbered as in the viral genome are shown at left. (B) Superposition of the degree of modification of each nucleotide on the best-fitting predicted secondary structure of MTE. Relative benzoyl cyanide modification is indicated by color-coding scheme of the SHAPE radioactivity scale. Nucleotides inside boxes are suspected to base pair with the 5’-UTR (Fig 7). Modification level of nucleotides in the presence of magnesium are indicated by stars following the reactivity color-coding scheme. Potential pseudoknot interaction between purine-rich bulge and bridging domain are indicated by dashed lines. *Note:* products of primer extension inhibition obtained from the SHAPE reaction are 1 nt shorter than those of dideoxy sequencing.

### Comparison of the MTE to PTEs: role of the pseudoknot

Because the MTE SHAPE probing experiments suggested that the MTE resembled a PTE, we compared the secondary structure of the MTE with known and predicted PTEs using the alignment program, LocARNA [54, 55] (Fig 4A). This alignment revealed a consensus structure with more variability than reported previously [53], because more predicted PTE sequences are aligned than previously. The MTE and PTEs contain a purine-rich bulge with at least one highly conserved G. However, the previously termed “C-rich” domain of PTE that bridges between stem-loops 1 and 2 is not always C-rich, thus we now call it the bridging domain. One putative PTE, from Pea stem necrosis virus (PSNV), contains no bridging domain and only a two base-pair stem in stem-loop 2 (Fig 4A). However, it has not been demonstrated to be functional. Potential pseudoknot base pairing between the purine-rich bulge and the bridging domain (square brackets Fig 4A), can be drawn for all PTEs except PSNV. However, for the MTE and some other PTE-like structures – the pseudoknot, if it exists at all, would consist of only two Watson-Crick base pairs: AG_4221-4222_:CU_4249-4250_ in the MTE. The SHAPE probing (Fig 3A) indicates that modification of AG_4221-4222_ decreased in the presence of Mg^2+^, and they are thus probably base-paired (which favors pseudoknot formation), but the already-modest SHAPE sensitivity of bases CU_4249-4250_ in the bridging domain does not decrease in the presence of Mg^2+^, as would be expected if the proposed pseudoknot forms. However, the bridging domain of other PTEs in which this pseudoknot is likely, also shows little change in modification in the presence of Mg^2+^ [53]. Thus, as with other PTEs, although phylogenetic and structural data suggest this pseudoknot occurs, we cannot conclude this without doubt.

**Fig 4.**
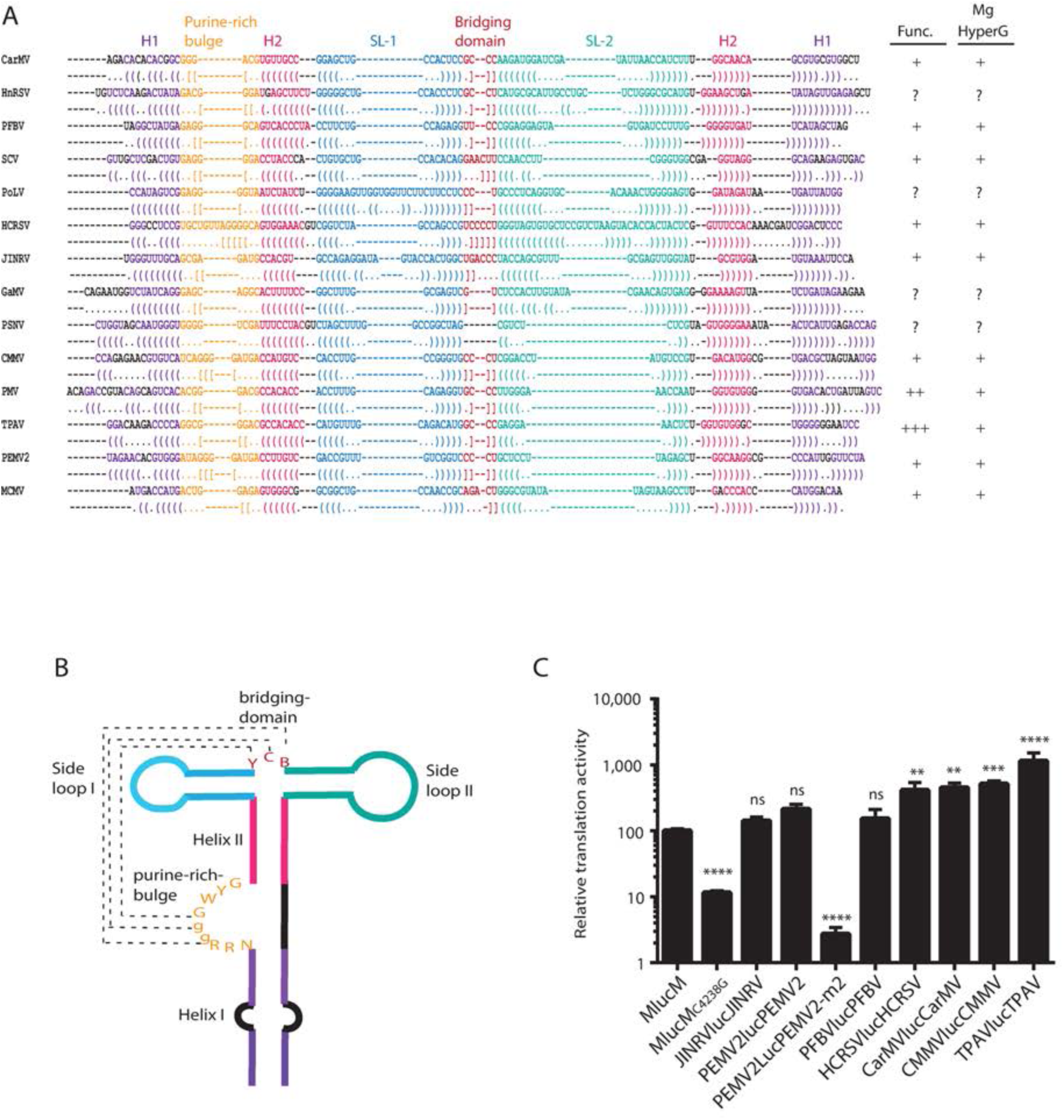
Alignment of known and predicted PTEs. (A) Secondary RNA structure alignment of PTEs. Structural input for alignment was used from previously published data (S2 Fig). LocARNA [55] was used to identify conserved regions of the PTE structure. Then structures were aligned to fit the consensus. Bases are color-coded based on the specific location in the PTE structure (panel B). Purple: conserved helix I region (H1), orange: conserved purine rich bulge, magenta: conserved helix II region (H2), blue: conserved stem-loop I (SL1), red: bridging domain, green: conserved stem-loop II (SL2). Parentheses or square brackets of same color in opposite orientation are below complementary bases. Square brackets show potential pseudoknot base pairing. (B) Sketch of conserved consensus shape of PTEs. Color-coding corresponds to alignment results on part (A). Conserved bases are shown using IUPAC nomenclature (Y = C or U, R = A or G, W = A or U, B = all except A). Lower case g indicates G in ≥78 % of PTEs. (C) Relative translation level activity of different PTEs in WGE with wild type MCMV indicated as 100%. Previously constructed luciferase reporter constructs contain a firefly luciferase reporter gene flanked by 5’ and 3’-UTRs of the indicated plant viral genomes containing PTE in the 3’ UTR [56]. PEMV2-m2 contains a CC to AA mutation in the bridging domain. Uncapped RNA transcripts were incubated for 30 min in WGE at room temperature. Data shown are percentage averages (±S.D.) from 3 independent experiments (n=9), with two, three and four asterisks indicating P<0.01, P<0.001, or P<0.0001, respectively, for significance of difference from wild type MCMV (MlucM) construct. ns = not significantly different from MCMV.

Because the MTE at least partially resembles the PTE consensus, we compared the MTE translation stimulation activity with that of other PTEs. Translation activity of luciferase reporter constructs containing PTEs in the 3’ UTR and the corresponding viral 5’ UTR [56] were compared to a construct containing MCMV 5’ and 3’ UTRs (MlucM). MlucM stimulated translation at a low level relative to most PTEs (Fig 4C). However, it stimulates translation 9-fold more than MCMV mutant C4238G, which prevents base pairing of the MTE to the 5’ UTR (below), and 20-fold more than the negative control PEMV2-m2, (a CC to AA mutation in the bridging domain), which was shown previously to virtually eliminate PTE activity [53, 56]. As reported previously [56], the PTE of Thin paspalum asymptomatic virus (TPAV) stimulated translation to a much higher level than the others. The PTEs of Japanese iris necrotic ring virus (JINRV) and Pelargonium flower break virus (PFBV) were not statistically significantly more stimulatory of translation than that of MCMV. Thus, even though the MTE resembles the PTE structure, it appears that MCMV (and JINRV and PFBV) have relatively weak PTE-like 3’ CITEs, compared to other characterized PTEs. It is noteworthy that here and previously [56], the most stimulatory PTEs (TPAV, PMV) have strong GGG:CCC pseudoknot base pairing between the purine-rich bulge and the bridging domain, whereas “weak” PTEs, such as the MTE and JINRV have little, if any pseudoknot base pairing (Fig 3B and S2 Fig, respectively). However, presence of a strong pseudoknot does not guarantee a strong translation enhancer, as indicated by PEMV2 and HCRSV PTEs (Fig 4A, Fig 4C, S1 Fig, S2 Fig). The role of potential pseudoknot base pairing is explored further below.

We constructed a series of mutations in the purine-rich bulge and bridging domain to test whether changes in these areas predicted to strengthen or weaken the pseudoknot had effects on translation efficiency (Fig 5). These included mutations designed to determine if a stronger pseudoknot could increase translation activity. Mutant A4248U, which should lengthen the proposed wild type pseudoknot from two (AG_4221-4222_:CU_4249-4250_) to three (AGA_4221-4223_:UCU_4248-4250_) base pairs, translated only 55% as efficiently as wild type in WGE (Fig 5B). This mutant could also potentially form an ACU_4216-4218_:AGU_4246-4248_ pseudoknot helix. To disrupt that possibility, a U4218A mutation was added. This double mutant translated 70% as efficiently as wild type in WGE (Fig 5C). However, neither of these mutants translated appreciably in the more competitive conditions in protoplasts. In other constructs, mutations in both the purine-rich bulge and the bridging domain were introduced to generate pseudoknot base pairing predicted to be more stable than wild type. In constructs in which the purine-rich bulge remained purine-rich and the bridging domain became pyrimidine-rich, changing the purine-rich domain or the bridging domain alone reduced translation (Fig 5 panels D, E, G, H), while the double mutants capable of forming the pseudoknot (GGG:CCC or AAAA:UUUU) translated more efficiently than the single-domain mutants. The GGG:CCC pseudoknot actually yielded 50% more luciferase than wild type in WGE and protoplasts (Fig. 5F), whereas the AAAA:UUUU predicted pseudoknot translated 35% as efficiently as wild type in WGE (Fig 5I), which was slightly greater than the UUU mutation alone (which may form a weak pseudoknot containing two G:U pairs) or the AAA mutation in the purine-rich bulge. Each of these mutants translated about 15-20% as efficiently as wild type in WGE. However, none of this set of mutants translated detectably in protoplasts (Fig 5, panels G, H, I). Swapping the purines and pyrimidines to create a potential: ACCC:GGGU pseudoknot helix gave low and no cap-independent translation in WGE and protoplasts, respectively (Fig 5J). One mutation, G4219U in the purine-rich bulge was not predicted to affect pseudoknot interactions and did not affect translation activity of the MTE (Fig 5K). This is interesting because G4219 is hypermodified in the presence of Mg^2+^ (Fig 3). Overall, with one rather modest exception (Fig 5F), mutations designed to increase pseudoknot base pairing altered the structure in such a way as to decrease translation efficiency.

**Fig 5.**
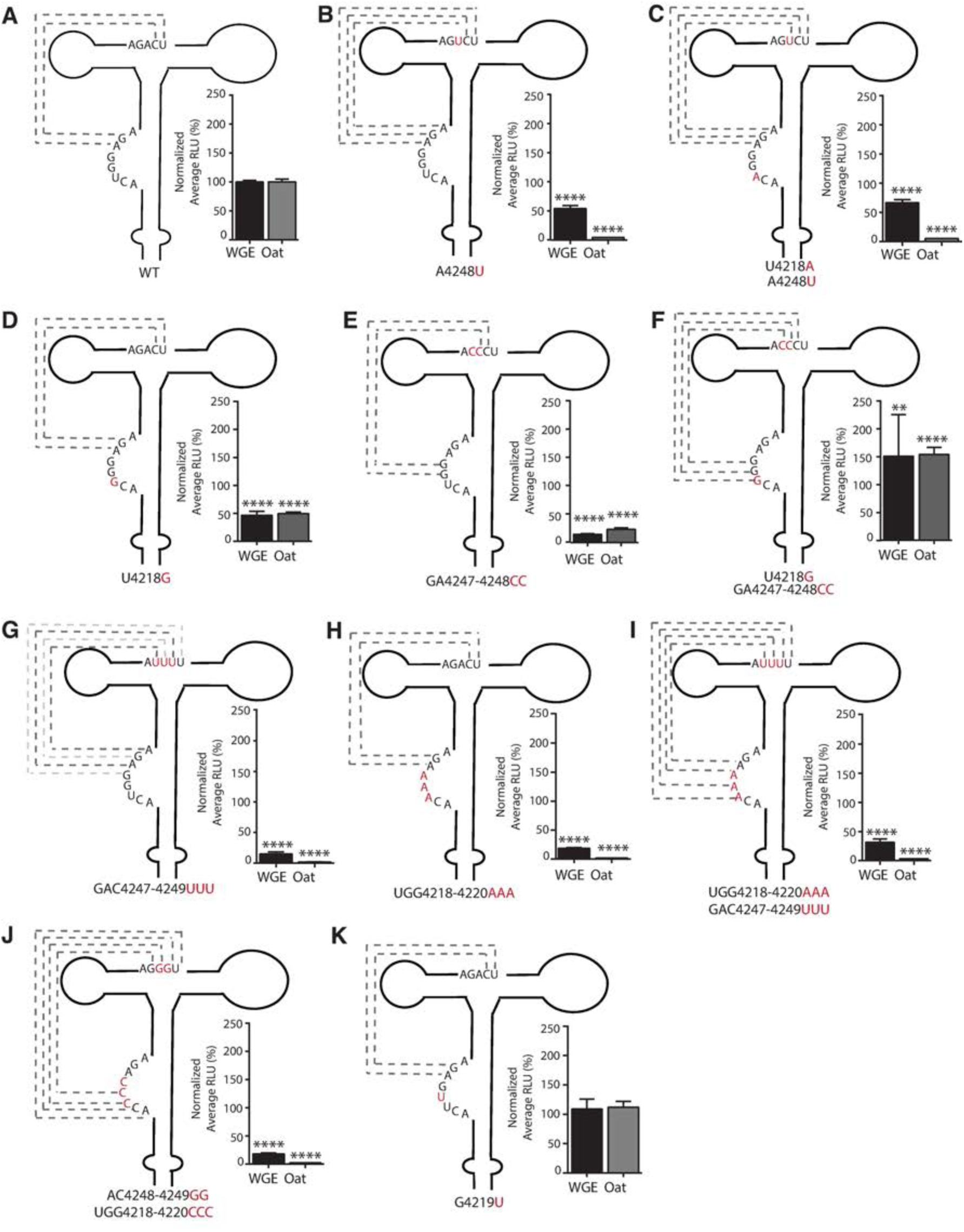
Effect of MTE mutations in the potential pseudoknot interaction on translation. Structures of predicted pseudoknot interaction mutants. Mutated bases in the MTE of each MlucM construct are in red. Constructs are named for the base changes at the numbered positions. Potential pseudoknot interactions are marked by dashed lines, with G:U pairs in lighter shade. Relative luciferase translation activities in WGE (black bars) or oat protoplasts (gray bars) are shown in percentages of relative light units normalized to MlucM wild type. Data are averages (±S.D.) from 4 independent experiments (n=15). Fluc/Rluc sample ratios were normalized to MlucM wt. One-way ANOVA was used to analyze the significance of each set of samples. One-way ANOVA-Dunnett’s multiple comparison test was performed to compare statistical difference among double mutants and single mutants. Four asterisks indicate statistical difference with P<0.0001; two asterisks indicate statistical difference with P<0.005

### The MTE binds eIF4E

Previously, PTEs have been shown to bind and require eukaryotic translation initiation factor 4E (eIF4E, the cap-binding protein), despite the absence of methylation (cap structure) on the PTE RNA [53, 57]. Because the MTE resembles PTEs, we used electrophoretic mobility shift assays (EMSA) to determine whether the MTE also binds eIF4E. To confirm eIF4E integrity (cap-binding ability), capped forms of the tested MTE constructs, were incubated in the presence of eIF4E and shown to confer strong mobility shifts (S3 Fig). We used the highly efficient TPAV PTE as a positive control for eIF4E binding to an uncapped PTE, as shown previously, and the nonfunctional TPAVm2, which contains mutations (CC to AA) in the bridging domain that inactivate the TPAV PTE and greatly reduced the binding affinity of the PTE to eIF4E (in the absence of a cap) as a negative control [56]. As expected, the TPAV PTE, formed a protein-RNA complex (Fig 6), as indicated by the reduced mobility of ^32^P-labeled PTE in the presence of eIF4E. Also as expected, the nonfunctional TPAVm2 PTE bound to eIF4E only at very high concentrations and most RNA remained unbound (Fig 6). Some nonspecific binding to any RNA by eIF4E is expected, as it is a low-affinity nonspecific RNA-binding protein [58, 59]. The wild type MTE_4187-4326_ migrated as two bands in the absence of added protein. Based on previous studies and migration patterns of other 3’ CITES [56, 57], we surmise that the fastest moving (and most abundant) band represents the properly folded MTE. Note that this band disappeared in the presence of ≥100 nM eIF4E (Fig 6B, MTE-wt). Thus, like PTEs, the MTE interacts with eIF4E with high affinity (Kd = 80-100 nM).

**Fig 6.**
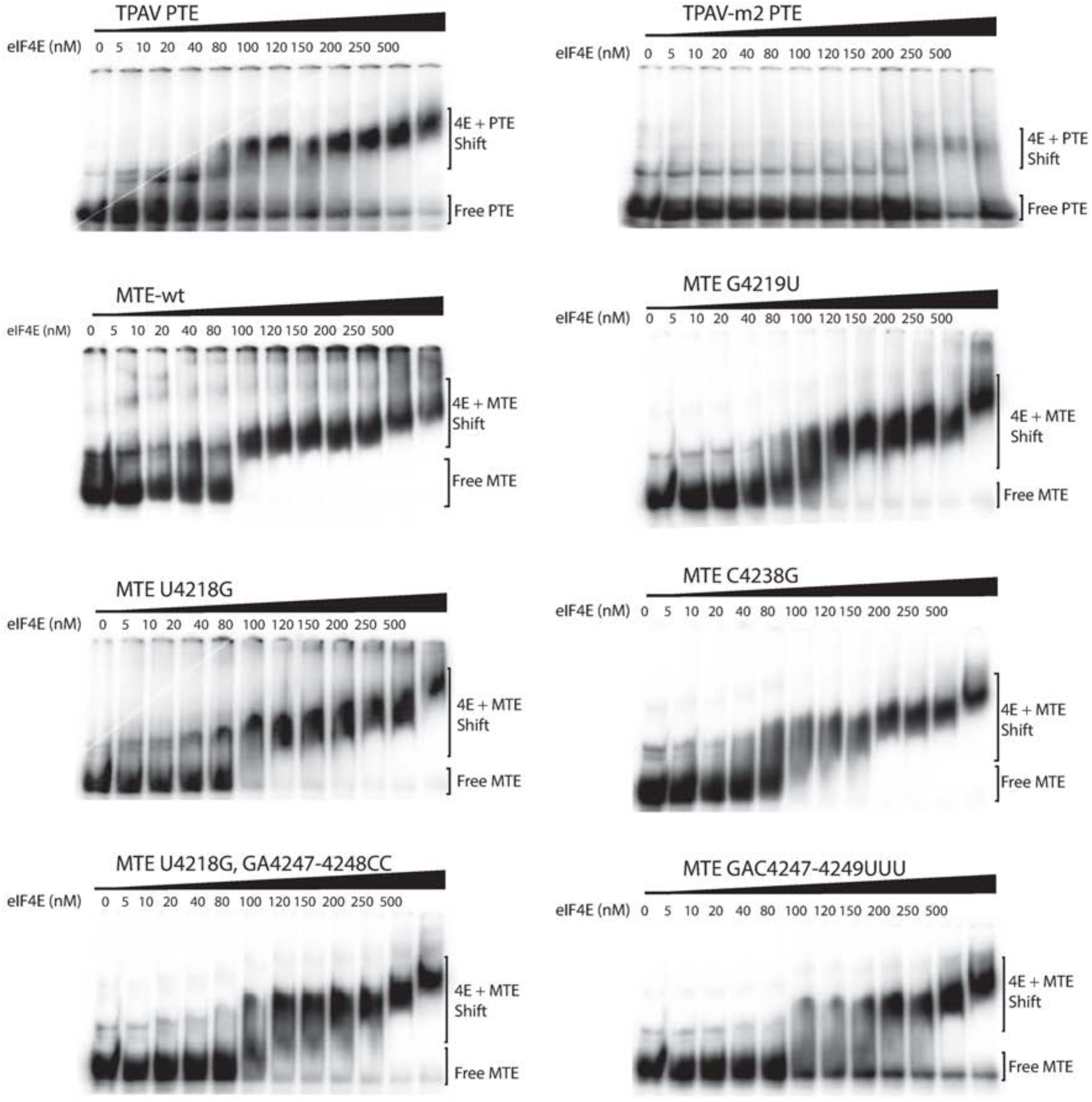
Electrophoretic mobility shift assays (EMSA) of mutant uncapped MTE and PTE RNAs with eIF4E. Ten fmol of the indicated uncapped ^32^P-labeled transcripts were incubated with the indicated concentrations of wheat eIF4E prior to electrophoresis on a non-denaturing gel.

We next tested the ability of mutant MTEs to bind eIF4E, in order to determine if eIF4E binding correlates with the translation enhancement function, as was observed previously for the TPAV PTE. Purine-rich bulge mutants U4218G and G4219U, which gave 50% and 100% of wild type translation, respectively, showed very similar EMSA profiles to wild type MTE (Fig 6). The mutant designed to strengthen the pseudoknot interaction, U4218G/GA4247-4248CC, and which gave 150% of wild type translation (Fig 5F) shifted similarly to U4218G alone, but with slightly less complete binding. This may be due to partial misfolding, as detected in the full-length genome context (below). MTE mutant GAC4247-4249UUU, which had greatly reduced translation, showed significantly less binding than the others at 80-200 nM eIF4E and some RNA remained unbound at the highest eIF4E concentrations. Finally, the mutant designed to prevent base pairing to the 5’ UTR (C4238G), but not expected to disrupt eIF4E binding because it enables cap-independent translation in the presence of complementary sequence in the 5’ UTR (shown below), gave an EMSA profile very similar to other functional mutants. In summary, functional MTEs bind eIF4E with higher affinity than a nonfunctional mutant, consistent with a requirement for eIF4E binding.

### Secondary structure of the MCMV 5’-UTR

Most plant viral 3’ CITEs that have been studied interact with the 5’ end of the viral genomic RNA and subgenomic mRNA via long-distance base pairing of the 3’ CITE to the 5’ UTR, presumably to deliver initiation factors to the 5’ end where they recruit the ribosomal 40S subunit to the RNA [35, 52, 60–62]. Thus, we sought to determine if the same interaction occurs in MCMV RNA. Initial *in silico* analysis of the 5’-end structure of MCMV revealed two sites (GGCA_12-15_ or UGGCA_103-107_) that could potentially base pair with the MTE sequence UGCCA_4236-4240_ in loop 1. An RNA transcript containing bases 1-140 of MCMV RNA, including the P32 (AUG_118-120_) and P50 (AUG_137-139_) ORF start codons, was subjected to SHAPE probing to determine which region was most likely to be available (single stranded) to interact with the MTE (Fig 7A). The 5’ UTR (nts 1-117) was found to consist of a large stem-loop with several large bulges, followed by a short stable stem-loop terminating 5 nt upstream of the start codon (Fig 7B). The first potential MTE-interacting sequence (GGCA_12-15_) is buried in a stem helix, while the UGGCA_103-107_ is in a favorable loop (Fig 7B). This led us to suspect that UGGCA_103-107_ is the potential base pairing sequence that interacts with MTE side loop 1.

**Fig 7.**
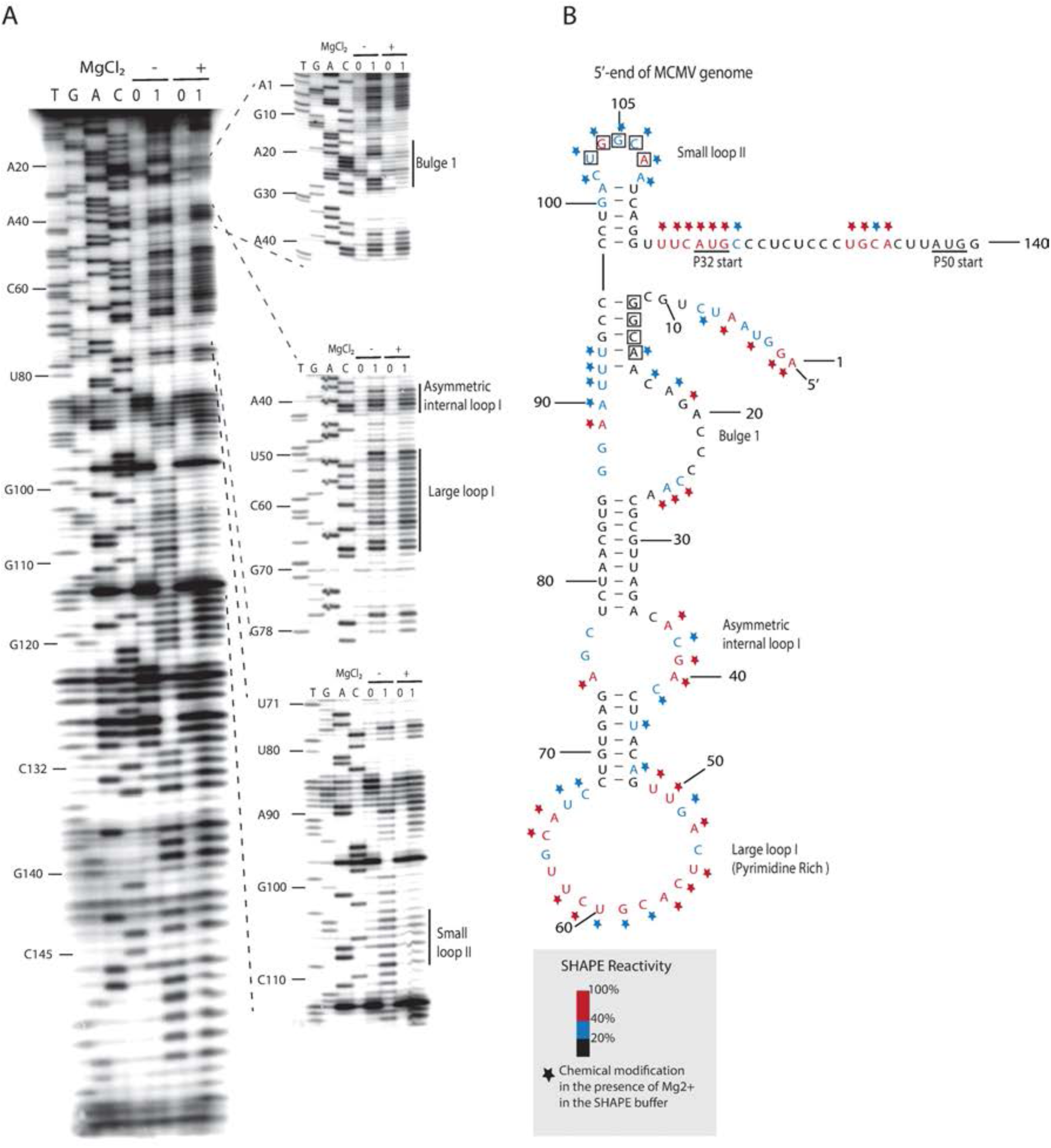
SHAPE analysis of the MCMV 5’ UTR. (A) SHAPE probing gel of the 5’ end of MCMV RNA (nts 1-140). RNA was treated with either 60 mM benzoyl cyanide (1) in dimethyl sulfoxide (DMSO) or DMSO only (0) in the presence (+) or absence (−) of magnesium in SHAPE buffer. The sequencing ladders (lanes TGAC) were generated by dideoxy sequencing of RNA with the same 5’-labeled primer used in the modification lanes. Zoom-in sections (gels at right) were obtained in a different gel by allowing electrophoresis to run longer. (B) Superposition of the probe activities on the best-fitting predicted secondary structure of the 5’ UTR. SHAPE activity level indicated by color-coded bases. Nucleotides inside the squares are tracts that have potential to base pair with loop I of the MTE (Fig 3). Modification level of nucleotides in the presence of magnesium are indicated by stars following the reactivity color-coding scheme. AUG start codons are underlined.

### Long distance base pairing of the MTE to the 5’ UTR

We next defined functionally which (if any) of the above candidate sequences base pairs to the MTE. Mutations were introduced in the XGCCA regions in the 5’ UTR (nts 11-15 or 103-107) and the MTE UGCCA_4236-4240_ region (Fig 8A). Mutation of G_13_ to C caused only a small decrease in luciferase activity in WGE and in oat protoplast translation systems (Fig 8B). In contrast, mutation of G_105_ to C, reduced luciferase activity by ∼75% in WGE and protoplasts. Even more extreme, the C4238G mutation of the middle base in the MTE UGCCA_4236-4240_ tract decreased luciferase activity by 80% to 90% (Fig 8B, see also Fig 4C). These mutations were then combined to restore any long-distance base pairing that may have been disrupted. The MlucM_G13C/C4238G_ double mutant, which would restore long-distance base pairing to the 5’-proximal complementary sequence in the 5’ UTR, yielded the same low translation activity as C4238G alone. In contrast, double mutant MlucM_G105C/C4238G_, which is predicted to restore long-distance base pairing of the MTE to the 5’ distal complementary sequence in the 5’ UTR, yielded a two-fold increase in translation activity when compared to MlucM_G105C_ and a 3-4-fold increase in translation relative to the more deleterious C4238G single mutation (Fig 8B). While the compensating mutations did not fully restore a wild type level of translation, the fact that the double mutant MlucM_G105C/C4238G_ but not MlucM_G13C/C4238G_ translated significantly more efficiently than MlucM_C4238G_ supports base pairing between 5’ UTR nts 103-107 and MTE nts 4236-4240 as a requirement for efficient cap-independent translation.

**Fig 8.**
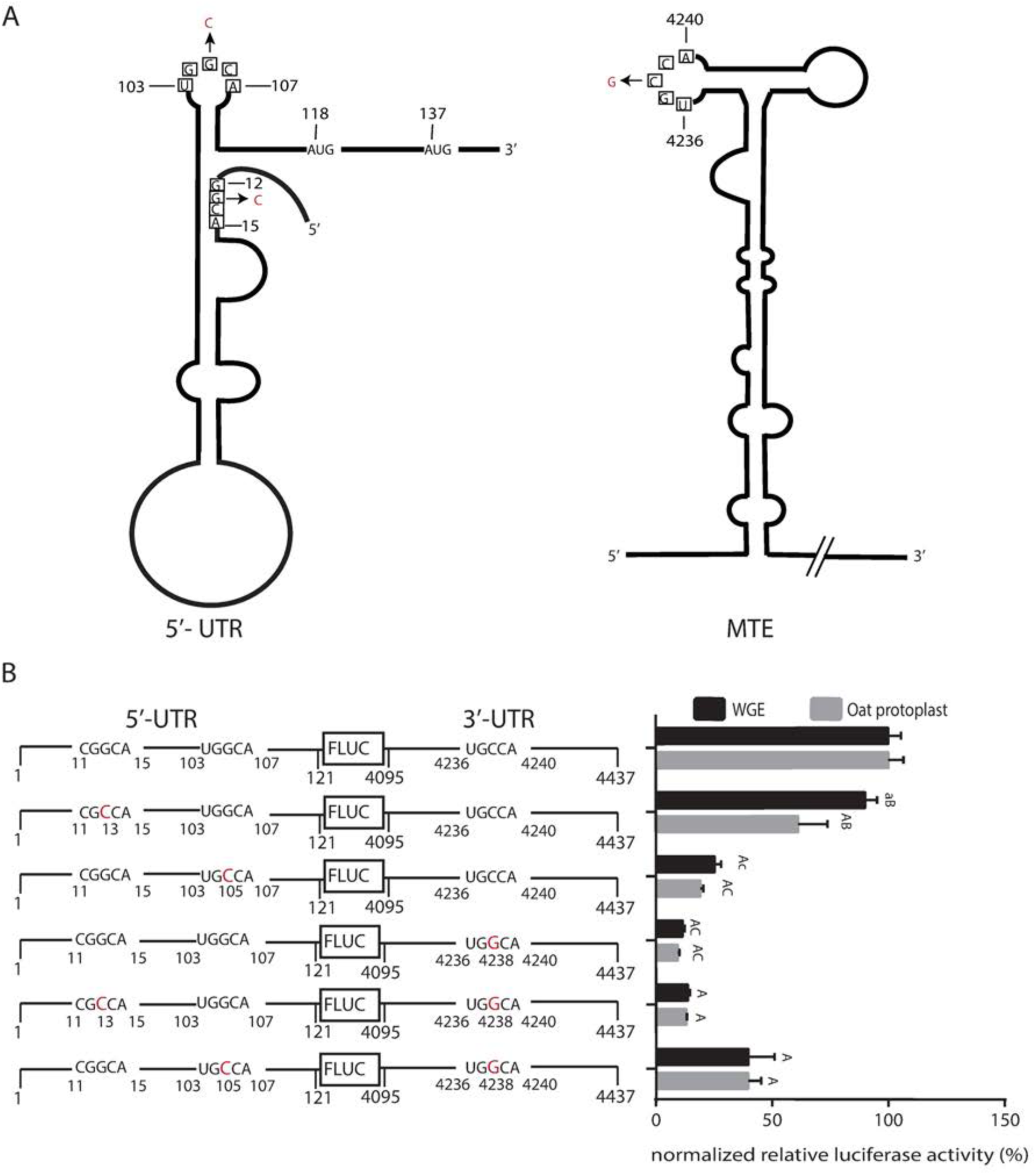
Long distance interaction between MCMV 5’ UTR and MTE. (A) Wire diagrams of 5’ UTR and MTE showing tracts (boxed bases) capable of base pairing between 5’ UTR and MTE. Mutations introduced to disrupt and restore potential long-distance base pairing are indicated in red. AUG start codons are shown at positions 118 and 137. (B) Effects of mutations (enlarged, red letters) on translation of MlucM. Relative luciferase translation activity of indicated uncapped transcripts in WGE and oat protoplasts are shown as percentages of relative light units relative to MlucM wild type (100%). Data are average percentages (±S.D.) from 4 independent experiments (for each construct: WGE: n=16; Oat: n=13). Two-way ANOVA multiple comparison was used to analyze the significance of each set of samples against MlucM_WT_ where A: P<0.0001, a: P<0.005. Two-way ANOVA Dunnett’s multiple comparison test was performed to compare statistical difference among double mutants and single mutants. Mutants compared with MlucM_G13C/C4238G_ were designated B if P<0.0001 or b if P<0.005. Mutants compared with MlucM_G105C/G4238G_ were designated C if P<0.0001 or c if P<0.005. ANOVA multiple comparisons were performed in the context of each sample collections (i.e. WGE or oat protoplast).

The MTE should also base pair to the 5’ UTR of sgRNA1, to allow translation of the viral coat and movement proteins. Indeed, we identified a sequence, UGGCA_2979-2983_ in the short 25 nt 5’ UTR of sgRNA1, which matches the UGGCA_103-107_ that base pairs to the MTE. This sequence is predicted to be in the terminal loop of the stem-loop that occupies the 5’ UTR of sgRNA1 (Fig 9A), which starts at nt 2971 [28]. We investigated both the effect of this short 25 nt 5’ UTR on MTE-mediated translation efficiency, and the role of base pairing (if any) between UGGCA_2979-2983_ and UGCCA_4236-4240_ in the MTE. In WGE, the sgRNA1 5’ UTR enabled translation about equally efficiently as did the genomic 5’ UTR, while translation was about two-thirds as efficient in oat protoplasts (Fig 9B), perhaps due to less RNA stability conferred by the shorter 5’ UTR. Separate G2981C and C4238G mutations in the sgRNA1 5’ UTR and the MTE, respectively, reduced translation significantly in both WGE and oat protoplasts. The double mutant, designed to restore predicted base pairing, with G2981C and C4238G in the same construct, gave surprising results. In WGE, as predicted, the double mutant fully restored translation to wild type levels. However, in oat protoplasts, the same mRNA was as nonfunctional as those containing single G2981C and C4238G mutations, showing no restoration of translation whatsoever. Because WGE is a high-fidelity translation system that measures only translation, independent of the complicated environment of the cell, we conclude that the long-distance base pairing is necessary for efficient cap-independent translation. We speculate that the G2981C mutation altered the RNA in such a way as to make it highly unstable in cells or able to fortuitously interact with cellular components that preclude translation, and which are absent in WGE.

**Fig 9.**
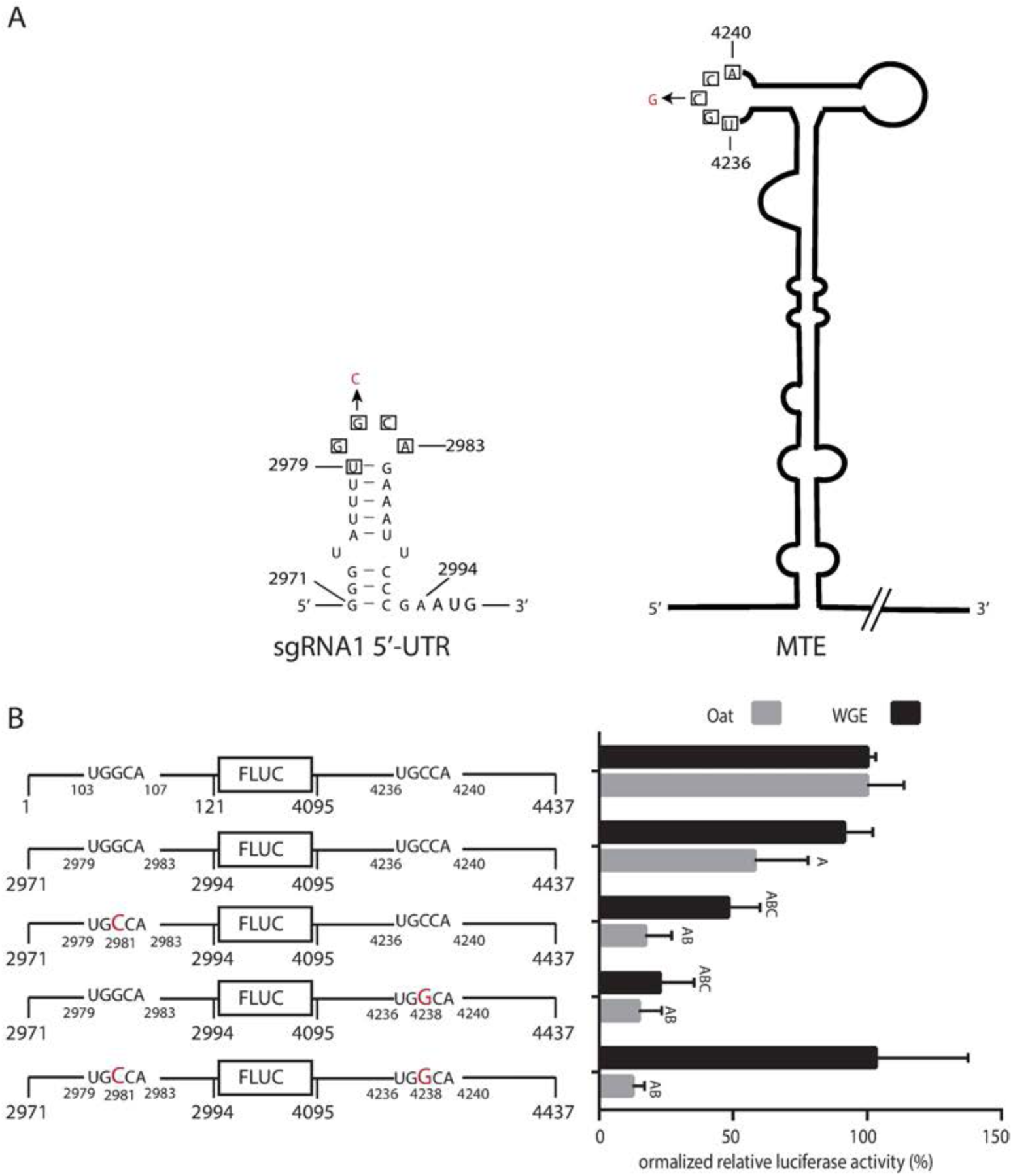
Long distance interaction between sgRNA1 5’ UTR and MTE. (A) Wire diagrams of sgRNA1 5’ UTR and MTE showing tracts (boxed bases) capable of base pairing between 5’ UTR and MTE. Mutations introduced to disrupt and restore potential long-distance base pairing are indicated in red. The first start codon is shown at nt 2995. (B) Translation of MlucM (top row) or Msg1lucM (remaining rows), with mutations shown in enlarged red text. Relative luciferase translation activity of indicated uncapped transcripts in WGE and oat protoplasts are shown as percentages of relative light units relative to MlucM wild type (100%). Data are average percentages (±S.D.) from 4 independent experiments (For each construct: WGE: n=12; Oat: n=16). One-way ANOVA multiple comparison was used to analyze the significance of each set of samples against MlucM_WT_ where A: P<0.0001. One-way ANOVA Dunnett’s multiple comparison test was performed to compare statistical difference among double mutants and single mutants. Mutants compared with Msg1lucM were designated B if P<0.0001. Mutants compared with Msg1lucM_G2981C/G4238G_ were designated C if P<0.0001. Mutations in UTRs are indicated in bold-red letters.

### Effects of mutations on translation of full-length MCMV genomic RNA

To determine the effects of mutations that affect translation in the natural context of genomic RNA, selected mutations from the luciferase experiments were introduced into the MCMV infectious clone pMCM41. Firstly, uncapped, full-length genomic RNA transcripts from pMCM41 mutants were translated in WGE, and the predominant ^35^S-met-labeled viral protein products (P50, P32 and P25) were observed. In agreement with the luciferase reporter constructs, mutants MCM41_C4238G_ and MCM41_G105C_, which disrupt the long-distance base pairing between MTE and 5’ UTR, yielded less viral protein than wild type (Fig 10A). In contrast, the double mutant, MCM41_G105C/C4238G_, which restores the long-distance base pairing, translated more efficiently than wild type RNA for the P32 and P50 proteins. A 25 kDa protein, presumably the viral coat protein (MW 25 kDa), was not expected to be translated much from genomic RNA as seen in Fig 1B, but it appeared in this experiment. Its translation remained at about the same reduced level in the double mutant as in the single mutants. MCM41 mutants G4219U, U4218G, and Δ4200-4300 translated similarly to the luciferase constructs, relative to wild type. However, the mutant designed to form a GGG:CCC pseudoknot in the MTE, MCM41_U4218G/GA4247-4248CC_, showed a substantial decrease in translation, in contrast to the same mutation in the luciferase reporter, in which translation was increased by 50% (Fig 5F).

**Fig 10.**
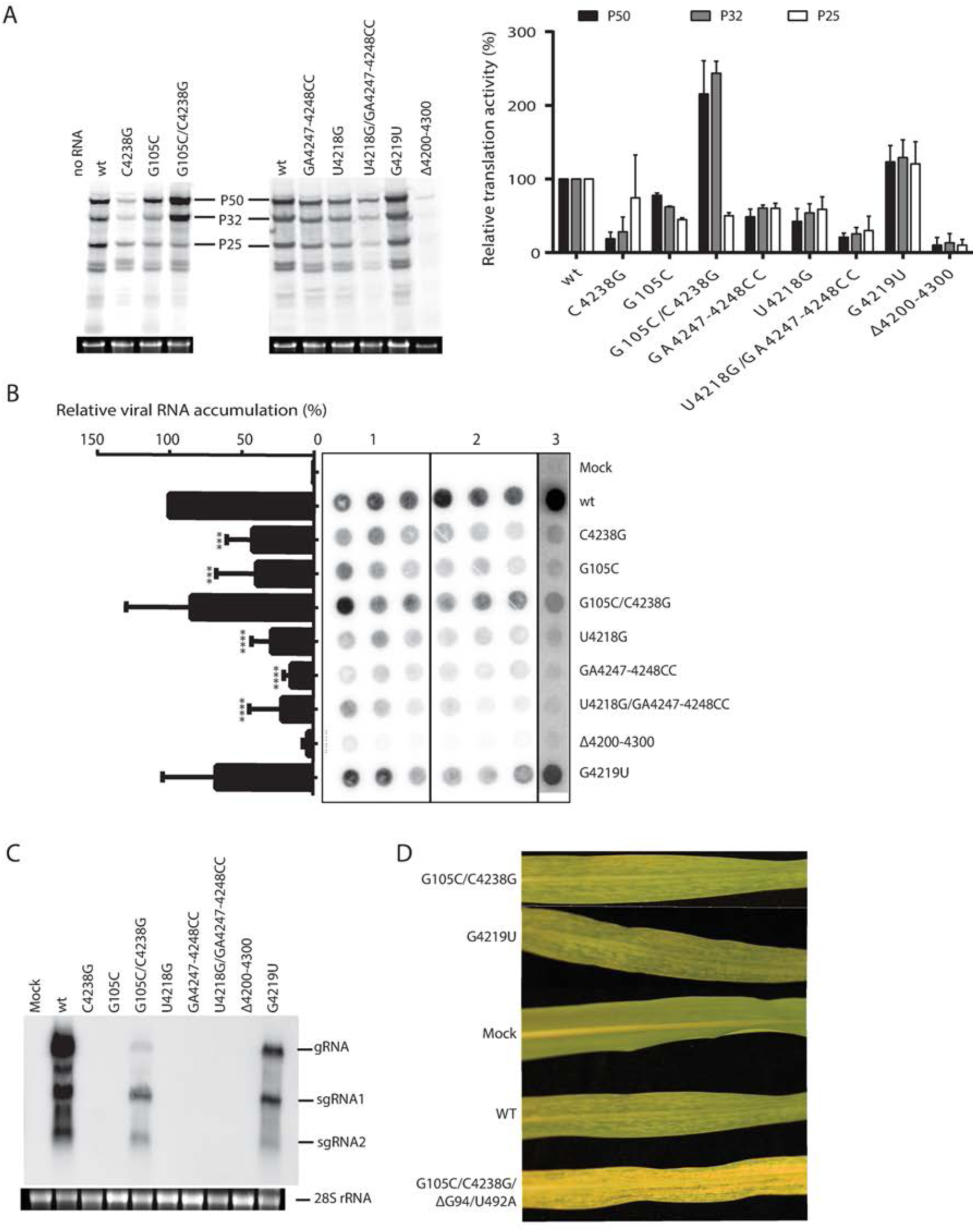
Effects of mutations on translation and replication of full-length MCMV genome. (A) Translation products from full-length MCM41 transcript in WGE. Prominent bands correspond to P50, P32, and P25 protein products. Percentage relative protein levels quantified using ImageQuant from two independent experiments was plotted in the graph at right. (B) Dot-blots of transfection assays in BMS protoplasts. 24 hr post transfection, total RNA was extracted and vacuum filtered through a nylon membrane (see Methods). Blots were probed with ^32^P-labeled antisense transcript complementary to nts 3811-4356 of MCMV genomic RNA, quantified by phosphorimagery, and normalized to that obtained with wild type MCM41 infection. One-way ANOVA was used to analyze the significance of each set of samples. Three, or four asterisks indicate statistical difference with P<0.001, or P<0.0001, respectively. Shown are blots from two experiments performed in triplicate, and one experiment with a single sample for each mutant. Mean relative spot intensity (viral RNA accumulation) for the three independent experiments was plotted in the bar graph to the left-side of the panel. (C) Northern blot hybridization of total RNA extracted from maize (B73) plants 14 dpi with the indicated mutants of MCM41. Mobilities of genomic RNA (gRNA), subgenomic RNAs 1 and 2 (sgRNA1 and sgRNA2) are indicated. (D) Symptoms at 14 dpi in systemically infected leaves rom plants inoculated with indicated mutants. The bottom leaf indicated the severe symptoms observed in plants infected with MCM41_G105C/C4238_ in which two spontaneous mutations, ΔG94 and U492A, appeared.

To determine if the mutants that translated poorly in the full-length genome context did so due to gross misfolding of the MTE, we performed SHAPE probing in the context of the MCMV genome (S6 Fig). Mutant C4238G, which disrupted long-distance base pairing with 5’UTR, maintained near-wild type MTE secondary structure, the only difference was the SHAPE reactivity data decreased in the side loop 1 (4235-4241) (S7 Fig). Functional mutants G4219U and U4218G were also similar in structure to WT (S7 Fig), with the exception that in U4218G, the purine-rich bulge was less modified in the absence and presence of magnesium (S6 Fig). Interestingly, secondary structures of the MTE mutants designed to have a strong pseudoknot, GA4247-4248CC and U4218G/GA4247-4248CC were changed radically, with either an increased SL-I stem at the expense of the pseudoknot, or formation of an unbranched, multiply-bulged stem-loop structure (S7 Fig). Both mutants changed the wild type MTE structure in such way that forced the reverse transcriptase to stop around nucleotides 4228-4241 (∼SLI) even in the absence of SHAPE chemicals (S6 Fig). These SHAPE results may explain the difference in function of this mutant between reporter assay (functional) and viral genomic context (nonfunctional).

### Replication of mutant MCMV RNA

To test the effects of mutations on viral RNA replication and accumulation, maize protoplasts from a Black Mexican Sweet (BMS) cell culture were transfected with full-length mutant MCM41 transcripts. Unfortunately, in BMS protoplast preparations, high levels of RNase obscured detection of distinct viral RNAs in northern blot hybridization, so the level of viral RNA was quantified by simple dot blot hybridization. Single mutants, which disrupted long distance base-pairing, MCM41_C4238G_ and MCM41_G105C_, yielded about 35-40% as much viral RNA replication product as wild type, while the double mutant, MCM41_G105C/C4238G,_ yielded about twice as much viral RNA as the single disruptive mutants (Fig 10B). On the other hand, the mutant designed to form a GGG:CCC pseudoknot, MCM41_U4218G/GA4247-4248CC_, yielded 80% less viral RNA than wild type, whereas replication of MCM41_G4219U_ was not significantly less than wild type. Finally, MCM41_Δ4200-4300_ produced virtually no viral RNA. Overall, efficiently translating mutants replicated at near-wild type levels, while mutants with reduced translation in the full-length context accumulated much lower levels of viral RNA, as expected.

We next attempted to validate the observations in protoplasts by inoculating maize plants with pMCM41 mutant transcripts (average of 36 individual inoculations per mutant). Maize B73 plants inoculated with wild type MCM41 transcript began to exhibit chlorotic mottling symptoms between 8-10 dpi, while sweet corn (Golden Bantam) plants exhibited symptoms around 6-9 dpi. Viral RNA was detected via RT-PCR in inoculated leaves of all plants at 8 dpi, including those that never showed chlorotic mottling (S5 Fig). At 14 dpi, samples from the newest systemic leaves were subjected to northern blot hybridization with a probe complementary to the 3’ end of the MCMV genome (Fig 10C). Following inoculation of both B73 and Golden Bantam maize plants, only three of the nine MCMV mutants tested elicited symptoms (chlorotic mottling). MCMV RNA was never detected in asymptomatic plants via northern blot hybridization. Viral RNA from samples displaying positive northern blot signals was subjected to Sanger sequencing for verification of introduced mutations (Table 1). The few single mutants (C4238G, G105C, U4218G) that showed symptoms at 14 dpi had reverted to wild type MCM41 (Table 1). MCM41_G4219U_ had a similar infectivity to wild type, but in about half of the infected plants the virus reverted to wild type (Table 1). Moreover, the MCM41_G4219U_ that did not revert to wild type accumulated less RNA (Fig. 10C), suggesting that, although the G4219U mutation is tolerated, the wild type sequence is more competitive. With one exception (below) MCM41_G105C/C4238G_ retained its introduced mutations but the infectivity (Table 1) and RNA accumulation (Fig 10C) also was reduced relative to wild type.

**Table 1.**
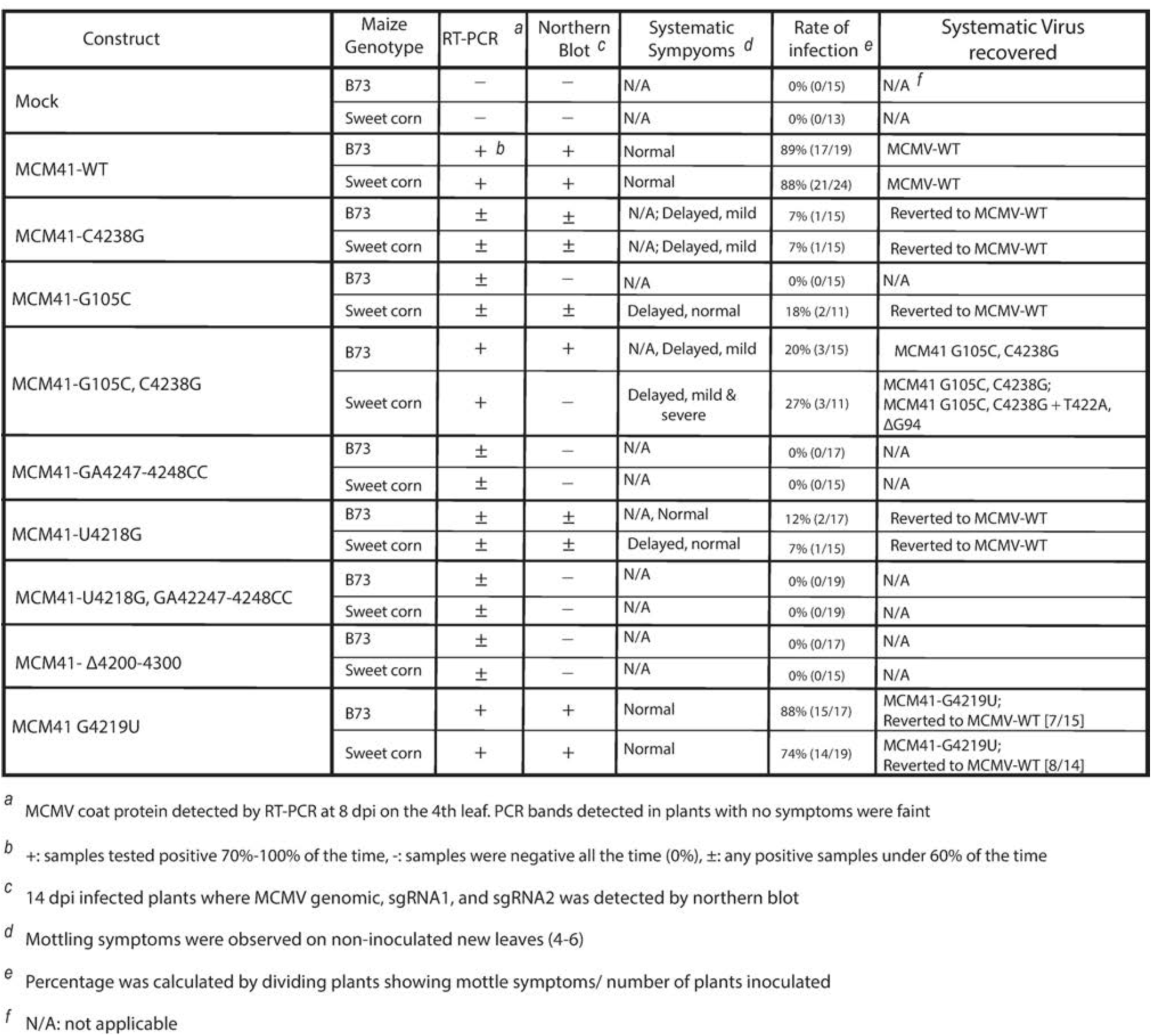

Interestingly, the viral RNA from one sweet corn plant inoculated with the MCM41_G105C/C4238G_ mutant acquired additional mutations. These spontaneous mutations consisted of deletion of G_94_ in the 5’ UTR, and a U492A point mutation in the overlapping P32 and P50 ORFs. This mutation introduced a stop codon in the P32 frame, shortening the protein from 289 aa to 125 aa, and changed amino acid 119 from valine to glutamic acid in the P50 and P111 (RdRp) proteins. This spontaneous mutant did not induce symptoms until 12 dpi, but after 14 dpi, symptoms were more extreme than wild type, giving nearly translucent leaves (Fig 10D). In summary, the MCMV genome tolerated few mutations, and only those mutations that allowed the most efficient translation replicated in maize plants.

## Discussion

### Identification of the 3’CITE in MCMV genome

Given the severe losses it has caused since 2011 in mixed infection with the potyvirus SCMV in East Africa [7, 63, 64], China [10, 65, 66] and Ecuador [13], and given the cost of screening seed worldwide to ensure absence of this seed-transmitted virus [63, 67, 68], MCMV is almost certainly the most economically important virus in the >76-member Tombusviridae family. Thus, it is imperative to understand the life cycle of MCMV, including translation, to identify molecular targets for genetic approaches to control this virus. All tombusvirids that have been studied contain a 3’ CITE [35, 40, 60, 69, 70], yet these are not predictable because the class of 3’ CITE a tombusvirid possesses does not always correlate with phylogeny [35, 52]. Moreover, we were unable to predict the class of 3’ CITE in MCMV RNA using ViennaFold or MFOLD. Therefore, we used experimental approaches to determine that MCMV contains a 3’ CITE in the PTE class, which we call MTE. Unlike the most efficient PTEs, the MTE lacks a strong pseudoknot and stimulates cap-independent translation less efficiently than most PTEs.

Previous studies of nine PTEs revealed a common secondary RNA structure composed of a branched structure with two side loops (Fig 4) connected by a pyrimidine-rich bridging domain [52] and a purine-rich bulge (formerly called G bulge) in the basal stem. A uniformly conserved feature is a guanosine in the purine-rich bulge that can be hypermodified by SHAPE reagents such as benzoyl cyanide in the presence of magnesium [53]. Despite this hypermodification, previous evidence also supports a pseudoknot interaction between purine-rich bulge bases and the bridging domain [53]. In the case of the MTE and some PTEs, the predicted pseudoknot interaction is tenuous, as the bridging domain is not always pyrimidine-rich, however, the magnesium-dependent hypermodified G in the G-bulge is still present. In the PMV and PEMV PTEs, this region is protected from SHAPE reagents by eIF4E and is thus the likely eIF4E-binding site [53]. The structural basis of this interaction is unclear, but the strongest PTEs have a relatively strong pseudoknot base pairing (GGG: CCC), whereas PTEs that have weak or no Watson-Crick base pairs are generally less stimulatory of translation (Fig 4). One mutant MTE designed to have a strong(er) pseudoknot yielded 50% more translation product than the wild type MTE in the luciferase reporter context (MlucM_U4218G/GA4247-4248CC_, Fig 5F), but these same mutations reduced translation, and thus replication, when in the context of the MCMV full-length clone (Fig 10) owing to misfolding (S7 Fig). Other mutants designed to have pseudoknots also reduced translation because of misfolding (S7 Fig).

### Long-distance kissing stem-loop interactions

A sequence in the 5’ UTR complementary to a loop in the 3’-CITE required for efficient translation has been found in most 3’-CITE-containing viral genomes [35]. This presumably facilitates initiation factor-mediated recruitment of the ribosomal subunit to the 5’-end of the RNA [35, 52, 60–62, 71].The sequence of the MTE long-distance kissing stem-loop interactions, between either the genomic 5’ UTR or the subgenomic RNA 5’ UTR and the MTE is UGGCA:UGCCA. This sequence fits the consensus found in many tombusvirids: YGGCA:UGCCR [35, 52], supporting our experimental evidence (Figs 8 and 9). Why this particular sequence pair has been conserved is unknown.

The unexpected translation of CP (P25) from full-length MCM41 RNA observed in Fig 10A, but only minimally in Fig 1B, may be due to different WGE batches. The two experiments were performed in Miller’s and Scheets’ labs, respectively. This may be due to variation in initiation factor levels or nucleases among batches of WGE used. It is possible that nucleolytic degradation in WGE generates small amounts of sgRNA1-sized RNA available for translation without internal ribosome entry. In fact, in WGE, the noncoding sgRNA2 is indeed generated by exonucleolytic degradation of all but the 3’ UTR of genomic RNA (our unpublished data, and [72]). Alternatively, internal ribosome entry is possible, as Simon’s lab has reported that another tombusvirid, TCV, harbors an internal ribosome entry site (IRES) to allow direct translation of CP from genomic RNA (as well as from sgRNA) [73]. However, the TCV IRES region is a tract of unstructured RNA, and we found no similar sequence upstream of the MCMV CP start codon.

The MCM41_C105G/G4238C_ transcript, which translates more efficiently than wild type in WGE, replicates indistinguishably from wild type MCM41 in protoplasts, but accumulates to much lower levels in whole plants. This is to be expected because the G4238C mutation in the MTE would prevent base pairing to the 5’ end of sgRNA1, which remains a wild type sequence. Thus, we predict translation of the CP and movement proteins is reduced in this construct. CP and movement proteins are not necessary for replication of other tombusvirids in protoplasts but would of course be necessary for the virus to accumulate in plants, thus explaining the difference in accumulation in Figs 10B and 10C.

### Translation of sgRNA1

Comparison of translation efficiencies of genomic (MlucM) and subgenomic RNA (Msg1lucM) reporter constructs revealed no striking difference in translation efficiency conferred by the 5’ UTR (Fig 9). This is interesting, because the secondary structure and length of the 5’ UTR is much less in the sgRNA1 5’ UTR (24 nt, ΔG = −6.6 kcal/mol) compared to that of genomic RNA 5’ UTR (117 nt, ΔG = −21.2 kcal/mol). We expected the sgRNA1 5’ UTR to provide superior translation efficiency because it should provide less resistance to ribosome scanning. Perhaps in infected cells, in the presence of the abundant sgRNA2, which contains the MTE, the sgRNA1 5’ UTR could outcompete the genomic RNA 5’ UTR for limiting eIF4E as discussed below.

In addition to stimulating translation of genomic RNA and sgRNA1 in *cis*, the MTE may regulate translation initiation in *trans*. Like certain other tombusvirids, such as red clover necrotic mosaic virus (RCNMV) [74, 75], tobacco necrosis virus-D [76] and the flaviviruses [77], MCMV generates a noncoding sgRNA (sgRNA2) corresponding to most of the 3’ UTR [28]. These RNAs are generated by exonucleolytic degradation of the larger viral RNAs until the exonuclease reaches a blocking structure called xrRNA at which point it stops, leaving the sgRNA, the 337 nt sgRNA2 in the case of MCMV, intact [72]. Because this abundant RNA contains the MTE, we propose that it regulates viral and host translation by binding and sequestering the eIF4E subunit of eIF4F [78]. sgRNA2 of BYDV (related to the Tombusviridae) binds the eIF4G subunit of eIF4F and, as a result, inhibits translation of BYDV genomic RNA, while favoring translation of BYDV sgRNA1, which, like MCMV sgRNA1, codes for movement and coat proteins. This would free genomic RNA of ribosomes, making it available for replication, encapsidation, and cell-to-cell movement. The selective inhibition of gRNA was shown to be due to its highly structured 5’ UTR, which likely increases dependence on eIF4F relative to the unstructured 5’ UTR of sgRNA1 [79]. The same regulation may occur on MCMV RNA. Thus, the MTE may play an essential role in MCMV infection by acting in *trans*. The role of sgRNA2 may be relevant to flaviviruses, which also produce noncoding sgRNAs from the 3’ UTRs that bind a variety of host proteins [77, 80–82], and which can affect translation [83].

### Efficient replication vs efficient translation?

It is possible that the relatively weak stimulation of translation by the MTE, and the relatively inefficient translation of viral genomic RNA, relative to artificially capped genomic RNA (Fig 1B), in contrast to the extremely high accumulation of MCMV RNA in infected cells (which is often visible by staining of total RNA from infected cells), may reveal a fundamental principle in RNA virus replication strategy. We propose that MCMV sacrifices efficient translation in exchange for extremely efficient RNA synthesis. RNA synthesis is inhibited by the presence of translating ribosomes on the viral RNA [82, 84, 85]. Thus, MCMV genomic RNA may have evolved to translate “poorly” in order to make it available to the viral replicase for highly efficient RNA replication. Moreover, owing to its abundance, genomic RNA may simply not need to translate efficiently. Indeed, highly efficient translation of such an abundant RNA could overwhelm the ribosomes, preventing translation of host mRNAs, harming the cell, and thus the virus itself. The 5’ UTR of the unrelated wheat yellow mosaic bymovirus harbors an IRES in a dynamic equilibrium state, which also gives suboptimal translation [86], perhaps also to allow efficient replication. In contrast, BYDV RNA has a highly efficient 3’ CITE (a BTE) [87], but accumulates to much lower levels in cells (not detectable by staining of total RNA). BYDV may have adopted a strategy to accumulate low levels of RNA and virus particles, which are adapted to very efficient acquisition by aphids. This RNA would require much more efficient translation in order to compete with host mRNAs for ribosomes. Thus, viruses such as MCMV are “replication strategists”, going “all in” on RNA synthesis at the expense of translation efficiency, while others, such as BYDV are “translation strategists”, translating efficiently at a cost to RNA replication.

### Toward host resistance to MCMV

As mentioned above, understanding translation may lead to strategies for resistance. The MTE binds eIF4E via what must include different molecular contacts than the binding of a 5’ m^7^G cap to eIF4E via the cap-binding pocket [88]. Thus, it may be possible to identify mutants of eIF4E that lose the ability to bind the MTE but retain a functional cap-binding pocket to allow translation of host (capped) mRNAs. In melon, such a resistance mechanism has been identified against melon necrotic spot virus (MNSV). A point mutation in melon eIF4E confers recessive resistance to most strains of MNSV, while having no negative effect on melon agronomic performance [89, 90]. This mutation was shown to preclude translation of MNSV RNA (thus blocking infection) by preventing efficient binding of eIF4E to the MNSV 3’ CITE (an I-shaped structure) [91]. In the case of MCMV, in addition to traditional screening of maize genotypes for MCMV resistance (which has achieved limited success), directed studies could identify natural eIF4E alleles, or guide construction of engineered eIF4E mutants, that bind poorly to the MTE, while still binding capped host mRNAs with high affinity, in order to achieve durable, recessive resistance to MCMV.

## Materials and methods

### Plasmids

The full length infectious clone of the MCMV genome (pMCM41) was described previously [29]. For psgRNA1 construction, template DNA pMCM41 was used to amplify MCMV nt 2972-4437 using Vent DNA polymerase, and oligonucleotides 3’SmaI (5’-agcaagcttcccGGGCCGGAAGAG [29] and sgRNA1 (5’GGTATTTTGGCAGAAATTCC) that were phosphorylated with T4 polynucleotide kinase. The vector pT7E19 [92] was digested with Sac I followed by mung bean nuclease digestion. Vector and insert were digested with Hind III, phenol/chloroform extracted, and precipitated prior to ligation with T4 DNA ligase. DNA was added to competent *E. coli* DH5α, selected on ampicillin/XGal plates, and screened by restriction digests and sequencing. Transcripts made from psgRNA1 linearized with SmaI contain MCMV nt 2971-4437, the complete sgRNA1. All enzymes were from New England Biolabs. MlucM was constructed using Gibson Assembly kit (New England Biolabs) such that a firefly luciferase (luc2, Promega) reporter gene was flanked by the 5’ and 3’ UTRs of MCMV (Fig 1). A Q5 Site-directed mutagenesis kit (New England Biolabs) with custom forward and reverse primers (S1 Table) was used to generate the deletions (Fig 2) and mutants (Figs 5, 8, 9) on the UTRs of the luciferase constructs. Resulting luciferase plasmid constructs were verified by sequencing at the Iowa State University DNA Sequencing Facility.

### *In vitro* Transcription

At Iowa State University (all experiments except Fig 1), plasmid DNA templates were linearized by restriction digestion or amplified by PCR to ensure correct template length. The RNAs were synthesized by *in vitro* transcription with T7 polymerase using MEGAscript (for uncapped RNAs) or mMESSAGE mMACHINE (for capped RNAs) kits (Ambion). RNAs used as probes for uncapped EMSA assays were generated using MEGAshortscript kit (Ambion). RNA transcripts were purified according to manufacturer’s instructions, RNA clean and concentrator kit (Zymo Research) was used for non-radioactive RNA preparation. RNase-free Bio-Spin columns P-30 (Bio-Rad) were used for radiolabeled RNA. RNA integrity was verified by 0.8% agarose gel electrophoresis. RNA concentration was determined by spectrophotometry for non-radiolabeled RNA. Radiolabeled RNA concentration was calculated by measuring the amount of incorporated radioisotope using a scintillation counter. At Oklahoma State University (Fig 1) full length (nt 1-4437) or 3’-truncated (nt 1-4195) genomic RNA was synthesized with T7 RNA polymerase (New England Biolab) and either pMCM41 or pMCM721 linearized with Sma I or Spe I following company protocols for uncapped or capped RNA synthesis. Unincorporated NTPs were removed by three rounds of ammonium acetate/ethanol precipitation and resuspension in nuclease-free water. pMCM721 contains a G residue between the T7 promoter and MCMV sequence, allowing synthesis of capped or uncapped RNAs 1 nt longer than WT [29]. The 3K MCMV RNA transcript templates were made by PCR using primers p9KO2 (CAGAAATTCCCGAgTGTC, nt 2982-2999) and M13 forward (−20) with both plasmid templates. These oligonucleotides were synthesized by the Oklahoma State University Core Facility.

### *In vitro* Translation

*In vitro* translation reactions were performed in WGE (Promega) as described [93]. Non-saturating amounts of RNAs (30 fmol) were translated in WGE in a total volume of 12.5 µl with amino acids mixture or [^35^S]-methionine amino acid mixture (C_t_= 3.06 μCi/0.14 MBq, 5.8 Ci/mmol, Perkin Elmer), 93 mM potassium acetate, and 2.1 mM MgCl_2_ based on manufacturer’s instructions. Translation reactions were incubated at room temperature (25°C) for 30 minutes. Translation products were separated by electrophoresis on a NuPAGE® 4-12% Bis-Tris gel (Invitrogen), detected with Pharox FX^TM^ plus Molecular Imager and quantified by Quantity One one-dimensional analysis software (Bio-Rad). For luciferase reactions, 10 µl of the translation reaction product was mixed into 50 µl of Luciferase assay reagent (Promega) and measured immediately on a GloMaxTM20/20 Luminometer (Promega). Statistical and data analysis were performed using GraphPad Prism software.

### Translation in Protoplasts

Uncapped MlucM (10 pmol) or its derived mutants were co-electroporated into ∼2×10^6^ oat (*Avena sativa* cv. Stout) protoplasts along with capped mRNA encoding *Renilla* luciferase (1 pmol) [94] as an internal control. Protoplasts were prepared and assays performed as described previously [95]. After 4 h of incubation at room temperature, protoplasts were harvested and lysed in Passive Lysis Buffer (Promega). Luciferase activities were measured using Dual Luciferase Reporter Assay System (Promega) in a GloMax^TM^ 20/20 luminometer (Promega). To minimized variation among electroporation replicates, firefly luciferase activities were normalized to the *Renilla* luciferase. Background, firefly relative light units (RLUs), measured in the absence of added luciferase mRNA, were subtracted from the values obtained with MlucM and its deletions/mutant derivatives. Statistical and data analysis was performed using GraphPad Prism software.

### RNA Structure Probing

Selective 2’-Hydroxyl Acylation analyzed by Primer Extension (SHAPE) was applied to selected UTR fragments of MCMV following the procedure previously described [96] except that SHAPE was conducted in the context of the complete genome instead of using the SHAPE cassette. In brief, 500 ng of RNA was denatured by heating to 95°C and renatured in SHAPE buffer (50mM HEPES-KOH, pH 7.2, 100 mM KCl, and ± 8 mM MgCl_2_) for 30 minutes at 30°C. RNA was modified by mixing 1/10 (v/v) ratio of renatured RNA with 60 mM benzoyl cyanide (Sigma) in anhydrous dimethyl sulfoxide (DMSO) (Sigma). After 2 min at room temperature, the RNA was mixed with four-fold excess tRNA and precipitated in 3 volumes of 99% ethanol and 1/10 volume 3 M sodium acetate. Control RNA was treated with the same amount of DMSO without benzoyl cyanide. The primer (3’UTR: 5’-TACTCCGTTGAGTTCAGAAACC-3’, or 5’UTR: 5’-CCATAAGTGCAGGGAGAGGG-3’) was 5’-end-labeled with γ-[^32^P] ATP and used for primer extension. Gel electrophoresis conditions and phosphorimager visualizations were done as described previously [57, 96].

### Expression and purification of wheat eIF4E

Wheat eIF4E pET22b plasmid clone [97] was obtained from Dr. Karen Browning. Plasmid was introduced in *E. coli* (BL21) cells and eIF4E expression was induced at O.D._600nm_= 0.7, with 0.5mM IPTG. From here on, the purification procedure of the protein was similar to previous published work [97]. In short, eIF4E expression in *E. coli* cells was induced by incubating in IPTG for 2 hours at 37°C with shaking (160 rpm). Cells were harvested by centrifugation at 6,000 x *g* for 15 minutes at 4°C, and cell pellets were quick frozen before purification. *E. coli* cells were disrupted by sonication in buffer B-50 (20 mM HEPES-KOH pH7.6, 0.1 mM EDTA, 1 mM DTT, 10% glycerol, 50 mM KCl) containing complete protease inhibitor cocktail tablets (Roche). Cell debris was separated from supernatant through 3-5 rounds of centrifugation (16, 000 x *g* for 15 minutes); for each centrifugation supernatant was transfer to a clean round-bottom centrifugation tube. Wheat eIF4E protein was purified using m^7^GTP agarose beads as described previously [97]. Protein was eluted using buffer B-100 (20 mM HEPES-KOH pH7.6, 0.1 mM EDTA, 1 mM DTT, 10% glycerol, 100 mM KCl) plus 20 mM of GTP. Protein purity was evaluated in a 4-12% NuPage Bis-Tris gel (Invitrogen). Protein concentration was determined by spectrophotometry and using Bio-Rad protein assay kit with a BSA protein standard curve.

### Electrophoretic mobility shift assay (EMSA)

Calculated specific activities of probes were used to determine the molarity of RNA in each purified stock. RNA probes were subjected to 6M urea-TBE gel electrophoresis to verify quality of RNA. RNA was diluted to 10 fmol/10μL for EMSA assays. As previously described [56],^32^P RNA labeled probes were incubated at indicated wheat eIF4E protein concentrations in EMSA binding buffer. Protein and RNA probes were incubated in a total volume of 10 μL of 1X EMSA binding buffer (10 mM HEPES pH 7.5, 20 mM KCl, 1 mM dithiothreitol, 3 mM MgCl_2_), 0.1 μg/μl yeast tRNA, 1 μg/μl bovine serum albumen, 1 unit/μl RNaseOUT ™ Recombinant Ribonuclease Inhibitor (Invitrogen), and 20mM Tris-HCl/10% glycerol for 25 minutes in ice. Then, 3 μL of 50% glycerol was added to each reaction. Immediately after, 7μL of RNA-protein mixture were loaded into a 5% polyacrylamide (acrylamide:*bis*-acrylamide 19:1), Tris-borate/EDTA (TBE) gel, which was run at ∼4°C at 110V for 45 minutes in 0.5X cold TBE buffer. Gels were dried on Whatman 3MM paper and exposed to a phosphorimager screen overnight. Phosphor screens were scanned in a Bio-Rad PhosphorImager, and radioactivity counts were analyzed using Quantity One software (Bio-Rad). Statistical analysis was performed using GraphPad Prism software.

### Inoculation of Black Mexican Sweet (BMS) protoplasts

Protoplasts were isolated from BMS suspension cultures as described previously [28, 29]. For each experiment, aliquots of isolated protoplasts (2.0 X 10^6^) were transfected with 5 pmol of pMCM41 and pMCM41 mutant transcript RNA using PEG-1540 (40% PEG, 6 mM CaCl_2_, 5 mM MES, pH 5.6). Protoplasts were diluted slowly in 5 mL of solution M (8.5% mannitol, 5 mM MES, pH5.6, 6 mM CaCl_2_), followed by incubation at 4°C for 20 minutes. Protoplasts then were centrifuged at 100 x *g* for 5 min, followed by two washes with solution M. PGM buffer (6% mannitol, 3% sucrose, M5524 MS salts (Sigma), M7150 vitamins (Sigma), 0.005% phytagel) was then added to each protoplast sample, followed by incubation in the dark at room temperature for 24 hours. RNA was isolated using Plant RNAeasy kit (Qiagen) following manufacturer’s instructions. Purified RNA was analyzed on a 0.8% native agarose gel and quantified using a Nanodrop Spectrophotometer.

### Plant Inoculations

Maize (B73 and sweet corn varieties) was grown in growth chambers on a 16:8 photoperiod with a temperature setting of 25°C/22°C (day/night). Maize seedlings were inoculated at the three-leaf stage. Plants were dusted with carborundum and inoculated on the third leaf with 10 μg of purified MCMV transcripts in 10mM sodium acetate, 5mM calcium chloride, and 0.5% bentonite by stroking three times with a freshly-gloved finger. Inoculated leaves were rinsed with water 10 minutes post inoculation; rinsed water was collected and autoclaved prior to disposal. Total plant RNA was extracted using Trizol (Invitrogen) from 100 mg of the newest systemic leaves for northern blot hybridization (14 dpi) and cDNA synthesis. Quality of RNA was evaluated in 0.8% native agarose gel, and RNA was quantified using a Nanodrop Spectrophotometer. RNA extracted from infected plants was subjected to cDNA synthesis and RT-PCR and sent to the Iowa State University DNA Facility for sequencing to evaluate the presence of introduced mutations.

### Dot blot and northern blot hybridizations

RNA isolated from BMS protoplasts had higher ratio of degraded RNA genomic/ sub-genomic RNA in northern blot hybridizations such that lower molecular weight fragments overpowered the genomics and sub-genomics RNAs. However, a trend on the amount of MCMV RNA detected suggested that replication was still occurring in protoplast; thus, dot blots were used instead. Unfractionated RNA from protoplasts was denatured prior to immobilization on a nylon membrane using a vacuum manifold apparatus as described in Brown et. al. 2004 [98]. RNA from maize plants was not degraded so it was subjected to northern blot hybridization. Total RNA from 100 mg newest leaf samples (14 dpi) was denatured in a formaldehyde/formamide buffer solution by heating at 65°C for 15 minutes, follow by separation in a 0.8% denaturing agarose gel and transferred to a nylon membrane. Both dot blot and northern blot nylon membranes were UV cross-linked, follow by prehybridization (50% formamide, 5X SSC, 200 μg/mL polyanetholesulfonic acid, 0.1% SDS, 20mM sodium phosphate, pH 6.5) at 55°C for 2 hours. Membranes were hybridized using ^32^P-labeled probe complementary to MCMV nts 3811-4356, which had been transcribed using SP6 RNA polymerase. Washed blots were wrapped in plastic and exposed to phosphor screens. Phosphor screens were scanned in a Bio-Rad PhosphorImager, radioactivity counts were analyzed using Quantity One software (Bio-Rad). Statistical analysis was performed using GraphPad Prism software.

### RT-PCR

Presence of MCMV virus RNA was evaluated by RT-PCR. Total RNA isolated from inoculated plants was extracted from washed maize 3^rd^ leaves using Trizol. The concentration and purity of extracted RNA was confirmed using a NanoDrop ND-2000 spectrophotometer and quality of RNA was evaluated via 0.8% native agarose gel electrophoresis. One microgram of RNA was subjected to cDNA synthesis following manufacturer’s instructions for Maxima first strand cDNA synthesis kit (Thermo Fisher Scientific). The cDNA was amplified using MCMV-CP primers (R: 5’-TGTGCTCAATGATTTGCCAGCCC, F: 5’-ATGGCGGCAAGTAGCCGGTCT) for 25 cycles, and the products were separated on 1% agarose gels, visualized by SYBR safe DNA stain (Thermo Fisher Scientific) and photographed. Similarly to MCMV-CP, maize *ubiquitin* cDNA expression (F: 5’-TAAGCTGCCGATGTGCCTGCGTCG and R: 5’-CTGAAAGACAGAACATAATGAGCACAG) was analyzed in the same sample set to serve as an endogenous positive control. Overall results from RT-PCR screenings were reported on Table 1. Examples of RT-PCR agarose gel observations can be observed on S5 Fig.

## Acknowledgements

The authors thank Dr. Karen Browning for advice on purification of eIF4E. Dr. Song Ki Cho and Dr. Jelena Kraft for advice on experimental protocols.

## Author Contributions

**Conceptualization:** W. Allen Miller

**Formal analysis:** Elizabeth J. Carino

**Investigation:** Elizabeth Carino, Kay Scheets

**Methodology:** Elizabeth Carino

**Supervision:** W. Allen Miller

**Writing – Original Draft Preparation:** Elizabeth J. Carino, Kay Scheets, W. Allen Miller

**Writing – Review & Editing:** Elizabeth J. Carino, W. Allen Miller

## Supporting information

**S1 Table. Table of primers.** List of primer sets used in this publication.

**S1 Fig. Luciferase translation assay including the 3’ end of coat protein coding region.** (A) Genome organization of MCMV. Boxes indicate open reading frames (ORFs) with encoded protein (named by its molecular weight in kDa) indicated. Black oval shows location of the 3’-CITE (4164-4333). Zigzag line indicates truncated sequence of the CP ORF included in luciferase construct. Luciferase translation assay results in WGE are shown to the right of the corresponding luciferase construct with the indicated 3’ UTR sequence. All constructs here contain the full MCMV 5’ UTR. Luciferase activities (relative light units) were normalized to MlucM construct containing the full 3’-UTR (4095-4437). Data samples were subjected to one-way ANOVA and Dunnett’s multiple comparison test was used to analyze the significance of each set of samples against MlucM where one asterisk equals 0.015 P-value. (B) *In vitro* translation assays using ^35^S-methionine. *Left*: a representative ^35^S-met-labeled gel sample out of 3 independent assays. *Right*: Quantification of the overall radioactivity detected in each assay.

**S2 Fig. Secondary structures of known and predicted PTEs.** The conserved bases predicted to interact with the 5’ UTR are inside black boxes. Predicted nucleotides which create the pseudoknot between the purine-rich bulge and the connecting bridge are indicated with dash lines.

**S3 Fig. Electrophoretic mobility shift assays (EMSA) of MTE and PTE capped RNA probes with eIF4E**. Ten fmol of the indicated capped ^32^P-labeled transcripts were incubated with the indicated concentrations of wheat eIF4E prior to electrophoresis on a non-denaturing gel.

**S4 Fig. Non-linear regression graphs of EMSAs.** Radioactivity in bands on gels in Fig 6 and S3 Fig was quantitated by phosphorimagery. The level of radioactivity in the unshifted, and shifted bands was used to generate nonlinear regression curves.

**S5 Fig. RT-PCR of plants inoculated with MCMV mutants.** Representative sample of results observed in RT-PCR screening of inoculated maize plants (8 dpi). RNA isolated from 3rd leaf was subjected to cDNA synthesis (see methods); cDNA from each isolation was divided into two fractions where one underwent RT-PCR to amplify Ubiquitin1 mRNA (top) as an internal control, while the other RT-PCR amplified the MCMV CP ORF (bottom). Summary of overall RT-PCR screenings is summarized on Table 1. For Table 1, even the faint bands observed in some mutants (e.g. samples 27-31) were counted as positive.

**S6 Fig. SHAPE probing of mutant MTE sequence in MCM41 mutant constructs used for infectivity assays**. (A) SHAPE probing gel of MTE in the whole genome context of MCM41 mutants. RNA was modified with either 60 mM benzoyl cyanide (1) in dimethyl sulfoxide (DMSO) or DMSO only (0) in the presence (+) or absence (−) of magnesium in SHAPE buffer. The sequencing ladders (lanes AC) were generated by dideoxy sequencing of RNA with the same 5’-labeled primer used in the modification lanes. (B) Reactivity information was superimposed into the characterized MTE structure. To compare where modifications occurred in the presence of magnesium among all mutants, specific geometric shapes were assigned to each hyper-modified nucleotide. Five-point star: wt, four-point star: C4238G, hexagon: GA4247-4248CC, rhombus: U4218G, triangle: GA4247-4248, U4218G, cross: G4219U. Color of each geometric figure corresponds to reactivity data coloring.

**S7 Fig. MTE SHAPE structures of selected mutants**. SHAPE data was superimposed into best fitted secondary structures. The base of the long stem on MTE was conserved throughout all the structures, thus only areas with major changes are shown. SHAPE reactivity data color-coding is the same as previous figure. Major changes in the MTE structure are indicated by the creamy-yellow shading.

## References

1. Niblett CL, Claflin LE. Corn lethal necrosis - a new virus disease of corn in Kansas. Plant Disease Reporter. 1978;62(1):15–9.

2. FAO. Maize Lethal Necrosis Disease (MLND) - A snapshot La FAO en situaciones de emergencias 2013 [March]. Available from: http://www.fao.org/fileadmin/user_upload/emergencies/docs/MLND%20Snapshot_FINAL.pdf.

3. FAO. Status of Maize lethal necrosis disease (MLND)in kenya Food and Agriculture Organization of the United Nations2017. Available from: https://www.ippc.int/en/countries/kenya/pestreports/2017/11/status-of-maize-lethal-necrosis-disease-mlndin-kenya-1/.

4. Wangai AW, Redinbaugh MG, Kinyua ZM, Miano DW, Leley PK, Kasina M, et al. First Report of Maize chlorotic mottle virus and Maize Lethal Necrosis in Kenya. Plant Dis. 2012;96(10):1582. Epub 2012/10/01. doi: 10.1094/PDIS-06-12-0576-PDN. PubMed PMID: 30727337.

5. Mahuku G, Lockhart BE, Wanjala B, Jones MW, Kimunye JN, Stewart LR, et al. Maize Lethal Necrosis (MLN), an Emerging Threat to Maize-Based Food Security in Sub-Saharan Africa. Phytopathology. 2015;105(7):956–65. Epub 2015/03/31. doi: 10.1094/PHYTO-12-14-0367-FI. PubMed PMID: 25822185.

6. Redinbaugh MG, Stewart LR. Maize Lethal Necrosis: An Emerging, Synergistic Viral Disease. Annu Rev Virol. 2018;5(1):301–22. Epub 2018/07/31. doi: 10.1146/annurev-virology-092917-043413. PubMed PMID: 30059641.

7. Gitonga K. Maize Lethal Necrosis - The growing challenge in Eastern Africa. In: Service UFA, editor. https://gain.fas.usda.gov: Global Agricultural Informaiton Network; 2014.

8. Kusia ES, Subramanian S, Nyasani JO, Khamis F, Villinger J, Ateka EM, et al. First Report of Lethal Necrosis Disease Associated With Co-Infection of Finger Millet With Maize chlorotic mottle virus and Sugarcane mosaic virus in Kenya. Plant Disease. 2015;99(6):899–900. doi: 10.1094/Pdis-10-14-1048-Pdn. PubMed PMID: WOS:000360866000056.

9. Wamaitha MJ, Nigam D, Maina S, Stomeo F, Wangai A, Njuguna JN, et al. Metagenomic analysis of viruses associated with maize lethal necrosis in Kenya. Virol J. 2018;15(1):90. Epub 2018/05/25. doi: 10.1186/s12985-018-0999-2. PubMed PMID: 29792207; PubMed Central PMCID: PMCPMC5966901.

10. Xie L, Zhang JZ, Wang QA, Meng CM, Hong JA, Zhou XP. Characterization of Maize Chlorotic Mottle Virus Associated with Maize Lethal Necrosis Disease in China. J Phytopathol. 2011;159(3):191–3. doi: 10.1111/j.1439-0434.2010.01745.x. PubMed PMID: WOS:000286838000011.

11. Deng TC, Chou CM, Chen CT, Tsai CH, Lin FC. First Report of Maize chlorotic mottle virus on Sweet Corn in Taiwan. Plant Dis. 2014;98(12):1748. Epub 2014/12/01. doi: 10.1094/PDIS-06-14-0568-PDN. PubMed PMID: 30703919.

12. Achon MA, Serrano L, Clemente-Orta G, Sossai S. First Report of Maize chlorotic mottle virus on a Perennial Host, Sorghum halepense, and Maize in Spain. Plant Disease. 2017;101(2):393-. doi: 10.1094/Pdis-09-16-1261-Pdn. PubMed PMID: WOS:000392355200054.

13. Quito-Avila DF, Alvarez RA, Mendoza AA. Occurrence of maize lethal necrosis in Ecuador: a disease without boundaries? European Journal of Plant Pathology. 2016;146(3):705–10. doi: 10.1007/s10658-016-0943-5.

14. Vega H, Beillard MJ. Ecuador declares state of emergency in corn production areas. In: Network GAI, editor. Quite, Ecuador: USDA Foreign Agricultural Service; 2016.

15. Goldberg KB, Brakke MK. Concentration of Maize Chlorotic Mottle Virus Increased in Mixed Infections with Maize-Dwarf Mosaic-Virus, Strain-B. Phytopathology. 1987;77(2):162–7. doi: DOI 10.1094/Phyto-77-162. PubMed PMID: WOS:A1987G522800008.

16. Scheets K. Maize chlorotic mottle machlomovirus and wheat streak mosaic rymovirus concentrations increase in the synergistic disease corn lethal necrosis. Virology. 1998;242(1):28–38. Epub 1998/03/17. doi: 10.1006/viro.1997.8989. PubMed PMID: 9501040.

17. Wang Q, Zhang C, Wang C, Qian Y, Li Z, Hong J, et al. Further characterization of Maize chlorotic mottle virus and its synergistic interaction with Sugarcane mosaic virus in maize. Sci Rep. 2017;7:39960. Epub 2017/01/07. doi: 10.1038/srep39960. PubMed PMID: 28059116; PubMed Central PMCID: PMCPMC5216416.

18. Xia Z, Zhao Z, Chen L, Li M, Zhou T, Deng C, et al. Synergistic infection of two viruses MCMV and SCMV increases the accumulations of both MCMV and MCMV-derived siRNAs in maize. Sci Rep. 2016;6:20520. Epub 2016/02/13. doi: 10.1038/srep20520. PubMed PMID: 26864602; PubMed Central PMCID: PMCPMC4808907.

19. Zambrano JL, Jones MW, Brenner E, Francis DM, Tomas A, Redinbaugh MG. Genetic analysis of resistance to six virus diseases in a multiple virus-resistant maize inbred line. Theor Appl Genet. 2014;127(4):867–80. Epub 2014/02/07. doi: 10.1007/s00122-014-2263-5. PubMed PMID: 24500307.

20. Gowda M, Beyene Y, Makumbi D, Semagn K, Olsen MS, Bright JM, et al. Discovery and validation of genomic regions associated with resistance to maize lethal necrosis in four biparental populations. Mol Breed. 2018;38(5):66. Epub 2018/05/19. doi: 10.1007/s11032-018-0829-7. PubMed PMID: 29773962; PubMed Central PMCID: PMCPMC5945787.

21. Redinbaugh MG, Zambrano JL. Chapter Eight - Control of Virus Diseases in Maize. In: Loebenstein G, Katis N, editors. Advances in Virus Research. 90: Academic Press; 2014. p. 391–429.

22. Xing Y, Ingvardsen C, Salomon R, Lubberstedt T. Analysis of sugarcane mosaic virus resistance in maize in an isogenic dihybrid crossing scheme and implications for breeding potyvirus-resistant maize hybrids. Genome. 2006;49(10):1274–82. Epub 2007/01/11. doi: 10.1139/g06-070. PubMed PMID: 17213909.

23. Jones MW, Penning BW, Jamann TM, Glaubitz JC, Romay C, Buckler ES, et al. Diverse Chromosomal Locations of Quantitative Trait Loci for Tolerance to Maize chlorotic mottle virus in Five Maize Populations. Phytopathology. 2018;108(6):748–58. Epub 2017/12/30. doi: 10.1094/PHYTO-09-17-0321-R. PubMed PMID: 29287150.

24. FAO. Central African Republic - Situation report March 2019 Food and Agriculture Organization of the United Nations2019 [cited 2019 March]. Available from: http://www.fao.org/fileadmin/user_upload/emergencies/docs/FAOCARsitrep_March2019.pdf.

25. FAO. Democratic Republic of the Congo - Situation report Februrary 2019 [Online report]. Food and Agriculture Organization of the United Nations 2019 [cited 2019 March]. Available from: http://www.fao.org/fileadmin/user_upload/emergencies/docs/FAO%20DRC%20sit%20rep_February%202019.pdf.

26. CIMMYT. Maize Lethal Necrosis: Building a comprehensive response. Report. Mexico: 2014.

27. King AM, Adams MJ, Carstens EB, Lefkowitz EJ. Machlomovirus. Virus Taxonomy. Report of the International Committee on Taxonomy of Viruses. 9th. United States of America: ELSEVIER Academic Press; 2012. p. 1134–8.

28. Scheets K. Maize chlorotic mottle machlomovirus expresses its coat protein from a 1.47-kb subgenomic RNA and makes a 0.34-kb subgenomic RNA. Virology. 2000;267(1):90–101. Epub 2000/01/29. doi: 10.1006/viro.1999.0107. PubMed PMID: 10648186.

29. Scheets K, Khosravi-Far R, Nutter RC. Transcripts of a maize chlorotic mottle virus cDNA clone replicate in maize protoplasts and infect maize plants. Virology. 1993;193(2):1006–9. Epub 1993/04/01. doi: 10.1006/viro.1993.1216. PubMed PMID: 8460472.

30. Nutter RC, Scheets K, Panganiban LC, Lommel SA. The complete nucleotide sequence of the maize chlorotic mottle virus genome. Nucleic Acids Res. 1989;17(8):3163–77. Epub 1989/04/25. doi: 10.1093/nar/17.8.3163. PubMed PMID: 2726455; PubMed Central PMCID: PMCPMC317721.

31. Scheets K. Analysis of gene functions in Maize chlorotic mottle virus. Virus Res. 2016;222:71–9. Epub 2016/06/01. doi: 10.1016/j.virusres.2016.04.024. PubMed PMID: 27242072.

32. Browning KS, Bailey-Serres J. Mechanism of cytoplasmic mRNA translation. Arabidopsis Book. 2015;13(13):e0176. Epub 2015/05/29. doi: 10.1199/tab.0176. PubMed PMID: 26019692; PubMed Central PMCID: PMCPMC4441251.

33. Browning KS. The plant translational apparatus. Plant Mol Biol. 1996;32(1-2):107–44. Epub 1996/10/01. doi: 10.1007/bf00039380. PubMed PMID: 8980477.

34. Walsh D, Mathews MB, Mohr I. Tinkering with translation: protein synthesis in virus-infected cells. Cold Spring Harb Perspect Biol. 2013;5(1):a012351. Epub 2012/12/05. doi: 10.1101/cshperspect.a012351. PubMed PMID: 23209131; PubMed Central PMCID: PMCPMC3579402.

35. Simon AE, Miller WA. 3’ cap-independent translation enhancers of plant viruses. Annu Rev Microbiol. 2013;67:21–42. Epub 2013/05/21. doi: 10.1146/annurev-micro-092412-155609. PubMed PMID: 23682606; PubMed Central PMCID: PMCPMC4034384.

36. Dreher TW, Miller WA. Translational control in positive strand RNA plant viruses. Virology. 2006;344(1):185–97. Epub 2005/12/21. doi: 10.1016/j.virol.2005.09.031. PubMed PMID: 16364749; PubMed Central PMCID: PMCPMC1847782.

37. Guo L, Allen E, Miller WA. Structure and function of a cap-independent translation element that functions in either the 3’ or the 5’ untranslated region. Rna. 2000;6(12):1808–20. Epub 2001/01/06. doi: 10.1017/s1355838200001539. PubMed PMID: 11142380; PubMed Central PMCID: PMCPMC1370050.

38. Miller WA, Wang Z, Treder K. The amazing diversity of cap-independent translation elements in the 3’-untranslated regions of plant viral RNAs. Biochem Soc Trans. 2007;35(Pt 6):1629–33. Epub 2007/11/23. doi: 10.1042/BST0351629. PubMed PMID: 18031280; PubMed Central PMCID: PMCPMC3081161.

39. Nicholson BL, White KA. 3’ Cap-independent translation enhancers of positive-strand RNA plant viruses. Curr Opin Virol. 2011;1(5):373–80. Epub 2012/03/24. doi: 10.1016/j.coviro.2011.10.002. PubMed PMID: 22440838.

40. Nicholson BL, Wu B, Chevtchenko I, White KA. Tombusvirus recruitment of host translational machinery via the 3’ UTR. Rna. 2010;16(7):1402–19. Epub 2010/05/29. doi: 10.1261/rna.2135210. PubMed PMID: 20507975; PubMed Central PMCID: PMCPMC2885689.

41. Gazo BM, Murphy P, Gatchel JR, Browning KS. A novel interaction of Cap-binding protein complexes eukaryotic initiation factor (eIF) 4F and eIF(iso)4F with a region in the 3’-untranslated region of satellite tobacco necrosis virus. J Biol Chem. 2004;279(14):13584–92. Epub 2004/01/20. doi: 10.1074/jbc.M311361200. PubMed PMID: 14729906.

42. Treder K, Pettit Kneller EL, Allen EM, Wang Z, Browning KS, Miller WA. The 3ʹ cap-independent translation element of Barley yellow dwarf virus binds eIF4F via the eIF4G subunit to initiate translation. Rna. 2008;14(1):134–47. doi: 10.1261/rna.777308.

43. Miras M, Truniger V, Querol-Audi J, Aranda MA. Analysis of the interacting partners eIF4F and 3’-CITE required for Melon necrotic spot virus cap-independent translation. Mol Plant Pathol. 2017;18(5):635–48. Epub 2016/05/05. doi: 10.1111/mpp.12422. PubMed PMID: 27145354; PubMed Central PMCID: PMCPMC6638222.

44. Huang M, Koh DC, Weng LJ, Chang ML, Yap YK, Zhang L, et al. Complete nucleotide sequence and genome organization of hibiscus chlorotic ringspot virus, a new member of the genus Carmovirus: evidence for the presence and expression of two novel open reading frames. J Virol. 2000;74(7):3149–55. Epub 2000/03/09. doi: 10.1128/jvi.74.7.3149-3155.2000. PubMed PMID: 10708431; PubMed Central PMCID: PMCPMC111815.

45. Altenbach SB, Howell SH. In vitro translation products of turnip crinkle virus RNA. Virology. 1982;118(1):128–35. Epub 1982/04/15. doi: 10.1016/0042-6822(82)90326-9. PubMed PMID: 18635130.

46. Johnston JC, Rochon DM. Translation of cucumber necrosis virus RNA in vitro. J Gen Virol. 1990;71 (Pt 10)(10):2233–41. Epub 1990/10/01. doi: 10.1099/0022-1317-71-10-2233. PubMed PMID: 2230730.

47. Na H, Fabian MR, White KA. Conformational organization of the 3’ untranslated region in the tomato bushy stunt virus genome. Rna. 2006;12(12):2199–210. Epub 2006/11/02. doi: 10.1261/rna.238606. PubMed PMID: 17077273; PubMed Central PMCID: PMCPMC1664717.

48. Koev G, Liu S, Beckett R, Miller WA. The 3prime prime or minute-terminal structure required for replication of Barley yellow dwarf virus RNA contains an embedded 3prime prime or minute end. Virology. 2002;292(1):114–26. Epub 2002/03/07. doi: 10.1006/viro.2001.1268. PubMed PMID: 11878914.

49. Zhang J, Simon AE. Importance of sequence and structural elements within a viral replication repressor. Virology. 2005;333(2):301–15. Epub 2005/02/22. doi: 10.1016/j.virol.2004.12.015. PubMed PMID: 15721364.

50. Zuker M. Mfold web server for nucleic acid folding and hybridization prediction. Nucleic Acids Res. 2003;31(13):3406–15. Epub 2003/06/26. doi: 10.1093/nar/gkg595. PubMed PMID: 12824337; PubMed Central PMCID: PMCPMC169194.

51. Lorenz R, Bernhart SH, Honer Zu Siederdissen C, Tafer H, Flamm C, Stadler PF, et al. ViennaRNA Package 2.0. Algorithms Mol Biol. 2011;6(1):26. Epub 2011/11/26. doi: 10.1186/1748-7188-6-26. PubMed PMID: 22115189; PubMed Central PMCID: PMCPMC3319429.

52. Chattopadhyay M, Kuhlmann MM, Kumar K, Simon AE. Position of the kissing-loop interaction associated with PTE-type 3’CITEs can affect enhancement of cap-independent translation. Virology. 2014;458–459:43-52. Epub 2014/06/15. doi: 10.1016/j.virol.2014.03.027. PubMed PMID: 24928038; PubMed Central PMCID: PMCPMC4101382.

53. Wang Z, Parisien M, Scheets K, Miller WA. The cap-binding translation initiation factor, eIF4E, binds a pseudoknot in a viral cap-independent translation element. Structure. 2011;19(6):868-80. Epub 2011/06/08. doi: 10.1016/j.str.2011.03.013. PubMed PMID: 21645857; PubMed Central PMCID: PMCPMC3113551.

54. Raden M, Ali SM, Alkhnbashi OS, Busch A, Costa F, Davis JA, et al. Freiburg RNA tools: a central online resource for RNA-focused research and teaching. Nucleic Acids Res. 2018;46(W1):W25–W9. Epub 2018/05/23. doi: 10.1093/nar/gky329. PubMed PMID: 29788132; PubMed Central PMCID: PMCPMC6030932.

55. Will S, Joshi T, Hofacker IL, Stadler PF, Backofen R. LocARNA-P: accurate boundary prediction and improved detection of structural RNAs. Rna. 2012;18(5):900–14. Epub 2012/03/28. doi: 10.1261/rna.029041.111. PubMed PMID: 22450757; PubMed Central PMCID: PMCPMC3334699.

56. Kraft JJ, Peterson MS, Cho SK, Wang Z, Hui A, Rakotondrafara AM, et al. The 3’ Untranslated Region of a Plant Viral RNA Directs Efficient Cap-Independent Translation in Plant and Mammalian Systems. Pathogens. 2019;8(1):28. Epub 2019/03/03. doi: 10.3390/pathogens8010028. PubMed PMID: 30823456; PubMed Central PMCID: PMCPMC6471432.

57. Wang Z, Treder K, Miller WA. Structure of a viral cap-independent translation element that functions via high affinity binding to the eIF4E subunit of eIF4F. J Biol Chem. 2009;284(21):14189–202. Epub 2009/03/12. doi: 10.1074/jbc.M808841200. PubMed PMID: 19276085; PubMed Central PMCID: PMCPMC2682867.

58. Magee J, Warwicker J. Simulation of non-specific protein-mRNA interactions. Nucleic Acids Res. 2005;33(21):6694–9. Epub 2005/11/30. doi: 10.1093/nar/gki981. PubMed PMID: 16314302; PubMed Central PMCID: PMCPMC1297708.

59. Carberry SE, Friedland DE, Rhoads RE, Goss DJ. Binding of protein synthesis initiation factor 4E to oligoribonucleotides: effects of cap accessibility and secondary structure. Biochemistry. 1992;31(5):1427–32. Epub 1992/02/11. doi: 10.1021/bi00120a020. PubMed PMID: 1737000.

60. Fabian MR, White KA. 5’-3’ RNA-RNA interaction facilitates cap- and poly(A) tail-independent translation of tomato bushy stunt virus mrna: a potential common mechanism for tombusviridae. J Biol Chem. 2004;279(28):28862–72. Epub 2004/05/05. doi: 10.1074/jbc.M401272200. PubMed PMID: 15123633.

61. Chattopadhyay M, Shi K, Yuan X, Simon AE. Long-distance kissing loop interactions between a 3’ proximal Y-shaped structure and apical loops of 5’ hairpins enhance translation of Saguaro cactus virus. Virology. 2011;417(1):113–25. Epub 2011/06/15. doi: 10.1016/j.virol.2011.05.007. PubMed PMID: 21664637; PubMed Central PMCID: PMCPMC3152624.

62. Miller WA, White KA. Long-distance RNA-RNA interactions in plant virus gene expression and replication. Annu Rev Phytopathol. 2006;44:447–67. Epub 2006/05/18. doi: 10.1146/annurev.phyto.44.070505.143353. PubMed PMID: 16704356; PubMed Central PMCID: PMCPMC1894749.

63. Braidwood L, Quito-Avila DF, Cabanas D, Bressan A, Wangai A, Baulcombe DC. Maize chlorotic mottle virus exhibits low divergence between differentiated regional sub-populations. Sci Rep. 2018;8(1):1173. Epub 2018/01/21. doi: 10.1038/s41598-018-19607-4. PubMed PMID: 29352173; PubMed Central PMCID: PMCPMC5775324.

64. Fatma HK, Tileye F, Patrick AN. Insights of maize lethal necrotic disease: A major constraint to maize production in East Africa. African Journal of Microbiology Research. 2016;10(9):271–9. doi: 10.5897/ajmr2015.7534.

65. Huang J, Wen GS, Li MJ, Sun CC, Sun Y, Zhao MF, et al. First Report of Maize chlorotic mottle virus Naturally Infecting Sorghum and Coix Seed in China. Plant Disease. 2016;100(9):1955-. doi: 10.1094/Pdis-02-16-0251-Pdn. PubMed PMID: WOS:000381658700043.

66. Wang Q, Zhou XP, Wu JX. First Report of Maize chlorotic mottle virus Infecting Sugarcane (Saccharum officinarum). Plant Dis. 2014;98(4):572. Epub 2014/04/01. doi: 10.1094/PDIS-07-13-0727-PDN. PubMed PMID: 30708716.

67. Uyemoto JK, Bockelman DL, Claflin LE. Severe Outbreak of Corn Lethal Necrosis Disease in Kansas. Plant Disease. 1980;64(1):99–100. doi: Doi 10.1094/Pd-64-99. PubMed PMID: WOS:A1980JL74200028.

68. Jiang XQ, Meinke LJ, Wright RJ, Wilkinson DR, Campbell JE. Maize Chlorotic Mottle Virus in Hawaiian-Grown Maize - Vector Relations, Host Range and Associated Viruses. Crop Prot. 1992;11(3):248–54. doi: Doi 10.1016/0261-2194(92)90045-7. PubMed PMID: WOS:A1992HU12400009.

69. Gao F, Simon AE. Differential use of 3’CITEs by the subgenomic RNA of Pea enation mosaic virus 2. Virology. 2017;510:194–204. Epub 2017/07/28. doi: 10.1016/j.virol.2017.07.021. PubMed PMID: 28750323; PubMed Central PMCID: PMCPMC5891822.

70. Truniger V, Miras M, Aranda MA. Structural and Functional Diversity of Plant Virus 3’-Cap-Independent Translation Enhancers (3’-CITEs). Frontiers in plant science. 2017;8:2047. Epub 2017/12/15. doi: 10.3389/fpls.2017.02047. PubMed PMID: 29238357; PubMed Central PMCID: PMCPMC5712577.

71. Sharma SD, Kraft JJ, Miller WA, Goss DJ. Recruitment of the 40S ribosome subunit to the 3’-untranslated region (UTR) of a viral mRNA, via the eIF4 complex, facilitates cap-independent translation. J Biol Chem. 2015;290(18):11268–81. Epub 2015/03/21. doi: 10.1074/jbc.M115.645002. PubMed PMID: 25792742; PubMed Central PMCID: PMCPMC4416834.

72. Steckelberg AL, Vicens Q, Kieft JS. Exoribonuclease-Resistant RNAs Exist within both Coding and Noncoding Subgenomic RNAs. MBio. 2018;9(6):e02461–18. Epub 2018/12/20. doi: 10.1128/mBio.02461-18. PubMed PMID: 30563900; PubMed Central PMCID: PMCPMC6299227.

73. May J, Johnson P, Saleem H, Simon AE. A Sequence-Independent, Unstructured Internal Ribosome Entry Site Is Responsible for Internal Expression of the Coat Protein of Turnip Crinkle Virus. J Virol. 2017;91(8):e02421–16. Epub 2017/02/10. doi: 10.1128/JVI.02421-16. PubMed PMID: 28179526; PubMed Central PMCID: PMCPMC5375686.

74. Iwakawa HO, Mizumoto H, Nagano H, Imoto Y, Takigawa K, Sarawaneeyaruk S, et al. A viral noncoding RNA generated by cis-element-mediated protection against 5’->3’ RNA decay represses both cap-independent and cap-dependent translation. J Virol. 2008;82(20):10162–74. Epub 2008/08/15. doi: 10.1128/JVI.01027-08. PubMed PMID: 18701589; PubMed Central PMCID: PMCPMC2566255.

75. Basnayake VR, Sit TL, Lommel SA. The Red clover necrotic mosaic virus origin of assembly is delimited to the RNA-2 trans-activator. Virology. 2009;384(1):169–78. Epub 2008/12/09. doi: 10.1016/j.virol.2008.11.005. PubMed PMID: 19062064.

76. Offel SK, Coutts RHA. Location of the 5’ Termini of Tobacco Necrosis Virus Strain D Subgenomic mRNAs. J Phytopathol. 1996;144(1):13–7. doi: 10.1111/j.1439-0434.1996.tb01481.x.

77. Clarke BD, Roby JA, Slonchak A, Khromykh AA. Functional non-coding RNAs derived from the flavivirus 3’ untranslated region. Virus Res. 2015;206:53–61. Epub 2015/02/11. doi: 10.1016/j.virusres.2015.01.026. PubMed PMID: 25660582.

78. Miller WA, Jackson J, Feng Y. Cis- and trans-regulation of luteovirus gene expression by the 3’ end of the viral genome. Virus Res. 2015;206:37–45. Epub 2015/04/11. doi: 10.1016/j.virusres.2015.03.009. PubMed PMID: 25858272; PubMed Central PMCID: PMCPMC4722956.

79. Shen R, Rakotondrafara AM, Miller WA. trans regulation of cap-independent translation by a viral subgenomic RNA. J Virol. 2006;80(20):10045–54. Epub 2006/09/29. doi: 10.1128/JVI.00991-06. PubMed PMID: 17005682; PubMed Central PMCID: PMCPMC1617300.

80. Damas ND, Fossat N, Scheel TKH. Functional Interplay between RNA Viruses and Non-Coding RNA in Mammals. Noncoding RNA. 2019;5(1):7. Epub 2019/01/17. doi: 10.3390/ncrna5010007. PubMed PMID: 30646609; PubMed Central PMCID: PMCPMC6468702.

81. Michalski D, Ontiveros JG, Russo J, Charley PA, Anderson JR, Heck AM, et al. Zika virus noncoding sfRNAs sequester multiple host-derived RNA-binding proteins and modulate mRNA decay and splicing during infection. J Biol Chem. 2019;294(44):16282–96. Epub 2019/09/15. doi: 10.1074/jbc.RA119.009129. PubMed PMID: 31519749; PubMed Central PMCID: PMCPMC6827284.

82. Sanford TJ, Mears HV, Fajardo T, Locker N, Sweeney TR. Circularization of flavivirus genomic RNA inhibits de novo translation initiation. Nucleic Acids Res. 2019;47(18):9789–802. Epub 2019/08/09. doi: 10.1093/nar/gkz686. PubMed PMID: 31392996; PubMed Central PMCID: PMCPMC6765113.

83. Holden KL, Harris E. Enhancement of dengue virus translation: role of the 3’ untranslated region and the terminal 3’ stem-loop domain. Virology. 2004;329(1):119–33. Epub 2004/10/13. doi: 10.1016/j.virol.2004.08.004. PubMed PMID: 15476880.

84. Murray KE, Steil BP, Roberts AW, Barton DJ. Replication of poliovirus RNA with complete internal ribosome entry site deletions. J Virol. 2004;78(3):1393–402. Epub 2004/01/15. doi: 10.1128/jvi.78.3.1393-1402.2004. PubMed PMID: 14722294; PubMed Central PMCID: PMCPMC321374.

85. Herold J, Andino R. Poliovirus RNA replication requires genome circularization through a protein-protein bridge. Mol Cell. 2001;7(3):581–91. doi: Doi 10.1016/S1097-2765(01)00205-2. PubMed PMID: WOS:000167966600013.

86. Geng G, Yu C, Li X, Yuan X. A unique internal ribosome entry site representing a dynamic equilibrium state of RNA tertiary structure in the 5’-UTR of Wheat yellow mosaic virus RNA1. Nucleic Acids Res. 2019. Epub 2019/11/13. doi: 10.1093/nar/gkz1073. PubMed PMID: 31713626.

87. Fan Q, Treder K, Miller WA. Untranslated regions of diverse plant viral RNAs vary greatly in translation enhancement efficiency. BMC Biotechnol. 2012;12(1):22. Epub 2012/05/09. doi: 10.1186/1472-6750-12-22. PubMed PMID: 22559081; PubMed Central PMCID: PMCPMC3416697.

88. Monzingo AF, Dhaliwal S, Dutt-Chaudhuri A, Lyon A, Sadow JH, Hoffman DW, et al. The structure of eukaryotic translation initiation factor-4E from wheat reveals a novel disulfide bond. Plant Physiol. 2007;143(4):1504–18. Epub 2007/02/27. doi: 10.1104/pp.106.093146. PubMed PMID: 17322339; PubMed Central PMCID: PMCPMC1851841.

89. Nieto C, Piron F, Dalmais M, Marco CF, Moriones E, Gomez-Guillamon ML, et al. EcoTILLING for the identification of allelic variants of melon eIF4E, a factor that controls virus susceptibility. BMC Plant Biol. 2007;7(1):34. Epub 2007/06/23. doi: 10.1186/1471-2229-7-34. PubMed PMID: 17584936; PubMed Central PMCID: PMCPMC1914064.

90. Truniger V, Nieto C, Gonzalez-Ibeas D, Aranda M. Mechanism of plant eIF4E-mediated resistance against a Carmovirus (Tombusviridae): cap-independent translation of a viral RNA controlled in cis by an (a)virulence determinant. Plant J. 2008;56(5):716–27. Epub 2008/07/23. doi: 10.1111/j.1365-313X.2008.03630.x. PubMed PMID: 18643998.

91. Nieto C, Morales M, Orjeda G, Clepet C, Monfort A, Sturbois B, et al. An eIF4E allele confers resistance to an uncapped and non-polyadenylated RNA virus in melon. Plant J. 2006;48(3):452–62. Epub 2006/10/10. doi: 10.1111/j.1365-313X.2006.02885.x. PubMed PMID: 17026540.

92. Petty IT. A plasmid vector for cloning directly at the transcription initiation site of a bacteriophage T7 promoter. Nucleic Acids Res. 1988;16(17):8738. Epub 1988/09/12. doi: 10.1093/nar/16.17.8738. PubMed PMID: 3047690; PubMed Central PMCID: PMCPMC338616.

93. Wang Z, Kraft JJ, Hui AY, Miller WA. Structural plasticity of Barley yellow dwarf virus-like cap-independent translation elements in four genera of plant viral RNAs. Virology. 2010;402(1):177–86. Epub 2010/04/16. doi: 10.1016/j.virol.2010.03.025. PubMed PMID: 20392470; PubMed Central PMCID: PMCPMC2905845.

94. Shen R, Miller WA. The 3’ untranslated region of tobacco necrosis virus RNA contains a barley yellow dwarf virus-like cap-independent translation element. J Virol. 2004;78(9):4655–64. Epub 2004/04/14. doi: 10.1128/jvi.78.9.4655-4664.2004. PubMed PMID: 15078948; PubMed Central PMCID: PMCPMC387721.

95. Rakotondrafara AM, Jackson JR, Kneller EP, Miller WA. Preparation and Electroporation of Oat Protoplasts from Cell Suspension Culture. Current Protocols in Microbiology: John Wiley & Sons, Inc.; 2005.

96. Miras M, Sempere R, Kraft J, Miller W, Aranda M, Truniger V. Determination of the Secondary Structure of an RNA fragment in Solution: Selective 2’-Hydroxyl Acylation Analyzed by Primer Extension Assay (SHAPE). Bio-Protocol. 2015;5(2):e1386. doi: 10.21769/BioProtoc.1386.

97. Mayberry LK, Dennis MD, Leah Allen M, Ruud Nitka K, Murphy PA, Campbell L, et al. Expression and purification of recombinant wheat translation initiation factors eIF1, eIF1A, eIF4A, eIF4B, eIF4F, eIF(iso)4F, and eIF5. Methods Enzymol. 2007;430:397–408. Epub 2007/10/05. doi: 10.1016/S0076-6879(07)30015-3. PubMed PMID: 17913646.

98. Brown T, Mackey K, Du T. Analysis of RNA by northern and slot blot hybridization. Curr Protoc Mol Biol. 2004;Chapter 4(1):Unit 4 9. Epub 2008/02/12. doi: 10.1002/0471142727.mb0409s67. PubMed PMID: 18265351.

