## Supplemental Data for "The RNA of maize chlorotic mottle virus - the essential virus in maize lethal necrosis disease - is translated via a panicum mosaic virus-like cap-independent translation element"

| Constructs: Deletion or mutation | Forward Primer | Reverse Primer |
|---|---|---|
| Δ4200-4300 | 5'-ACGGTGCGACATGGTAAC-3' | 5'-AACTACTAGTATACGATTTAGGC-3' |
| Δ4095-4164 | 5'-AAAGTGTTTGGGAGCCTAAATC-3' | 5'-CGCGGCTTACAATTTGGACTTTC-3' |
| Δ4095-4191 | 5'-CTAGTAGTTTGCGTGATGAC-3' | 5'-CGCGGCTTACAATTTGGACTTTC-3' |
| AC4248-4249GG | 5'-GCCAACCGCAGGGTGGGCGTATATAGTAAGCCTTGACC-3' | 5'-AGCCGCCGCCCACTCTCC-3' |
| UGG4218-4220CCC | 5'-TGACCATGACCCCAGAGTGGGCG-3' | 5'-TCACGCAAACTACTAGTATAC-3' |
| GA4247-4248CC | 5'-GCCAACCGCACCCTGGGCGTATATAGTAAGCCTTGACCC-3' | 5'-AGCCGCCGCCCACTCTCC -3' |
| U4218G | 5'-TGACCATGACGGGAGAGTGGG-3' | 5'-TCACGCAAACTACTAGTATAC-3' |
| GAC4247-4249UUU | 5'-GCCAACCGCATTTTGGGCGTATATAGTAAGCCTTGACCCAC-3' | 5'-AGCCGCCGCCCACTCTCC -3' |
| UGG4218-4220AAA | 5'-TGACCATGACAAAAGAGTGGGCG -3' | 5'-TCACGCAAACTACTAGTATAC -3' |
| G4219U | 5'-TGACCATGACTtGAGAGTGGG-3' | 5'-TCACGCAAACTACTAGTATAC-3' |
| A4248U | 5'-GCCAACCGCAGTCTGGGCGTATATAGTAAGCCTTGACC-3' | 5'-AGCCGCCGCCCACTCTCC-3' |
| U4218A | 5'-TGACCATGACAGGAGAGTGGG-3' | 5'-TCACGCAAACTACTAGTATAC-3' |
| MTE EMSA probe | 5'-AATTAATACGCTCACTATAGGGTATACTAGTAGTTTGCGT-3' | 5'-TGTATCCAGTTACCATGTCGCACCG-3' |
| TPAV EMSA probe | 5'-taatacgactcactataggGCTCTATCCGAAACTCCAGTG-3' | 5'-ACTCTCTACCTTCCGTCCAG-3' |
| G13C | 5'-GTAATCTGCGCCAACAGACCC -3' | 5'- CTCCCTATAGTGAGTCGTATTAGTG -3' |
| G105C | 5'-CCCCTGACTGCCAATCAGGTTTC -3' | 5'-AAATTCCCACGTTAGAGCTC -3' |
| C4238G | 5'-GCGGCGGCTGgCAACCGCAGA-3' | 5'-CCACTCTCCAGTCATGGTCATCACGCAAAC-3' |
| sg1MlucM | 5'-AGAAATTCCCGACGCGCGCATGGAAGACGC -3' | 5'-GCCAAAATACCCCCTATAGTGAGTCGTATTAGTGGCTTTACC -3' |
| G2981C | 5'-AGAAATTCCCGACGCGCGCATGGAAGACGC -3' | 5'-GGCAAAATACCCCCTATAGTGAGTCGTATTAGTGGCTTTACC -3' |
| MCMV-coat protein | 5'-ATGGCGGCAAGTAGCCGGTCT -3' | 5'- TGTGCTCAATGATTTGCCAGCCC -3' |
| Maize Ubiquitin 1 | 5'-TAAGCTGCCGATGTGCCTGCGTCG -3' | 5'- CTGAAAGACAGAACATAATGAGCACAG -3' |
| Northern Blot probe | 5'-ctgaATTTAGGTGACACTATAGTTCCTAGCATCTACTTGCCCC -3' | 5'-GGATCGTGCCCTCAGCTACAATAGCTCTGAA -3' |

Supplementary Table S-T1

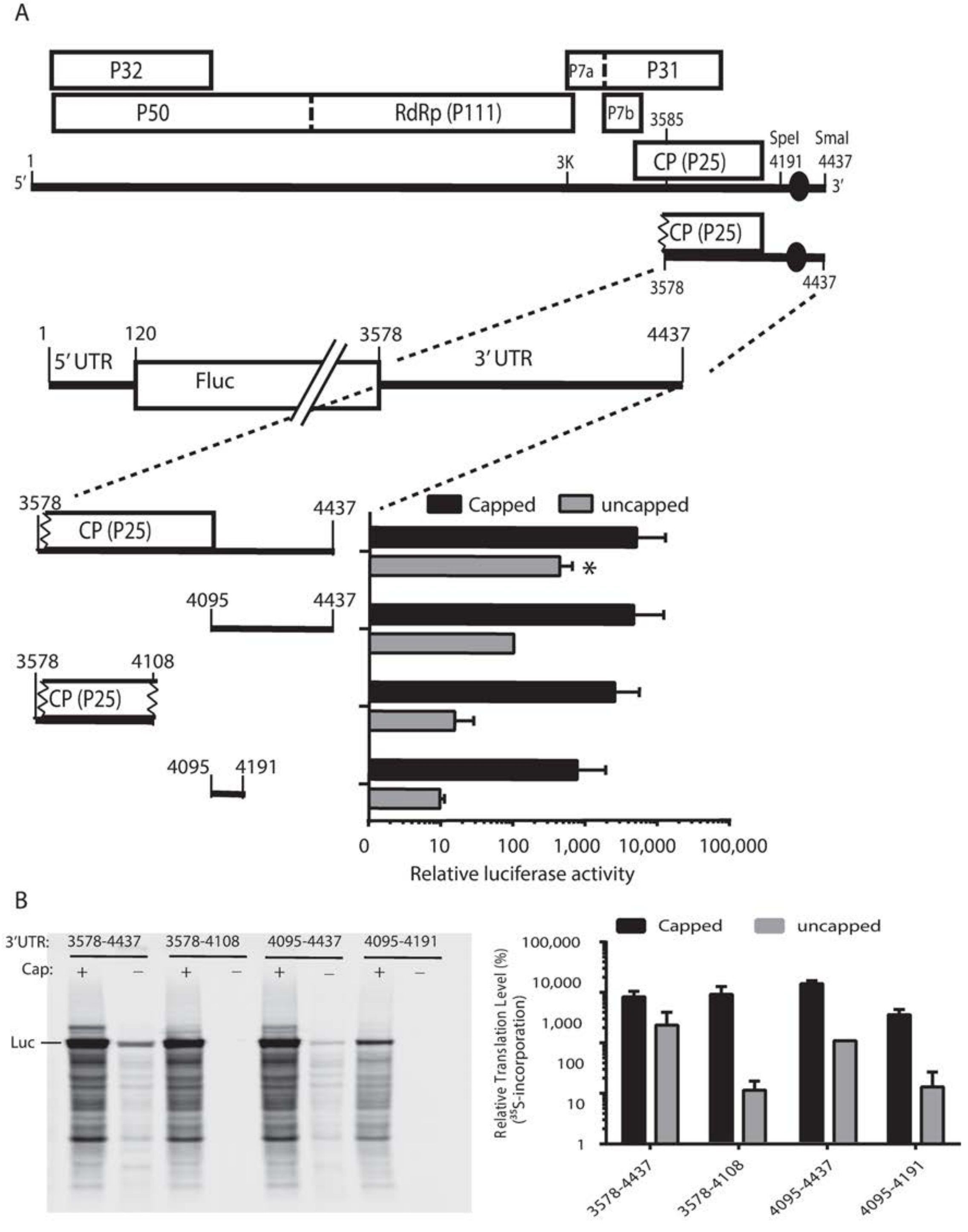


Supplementary Figure S1

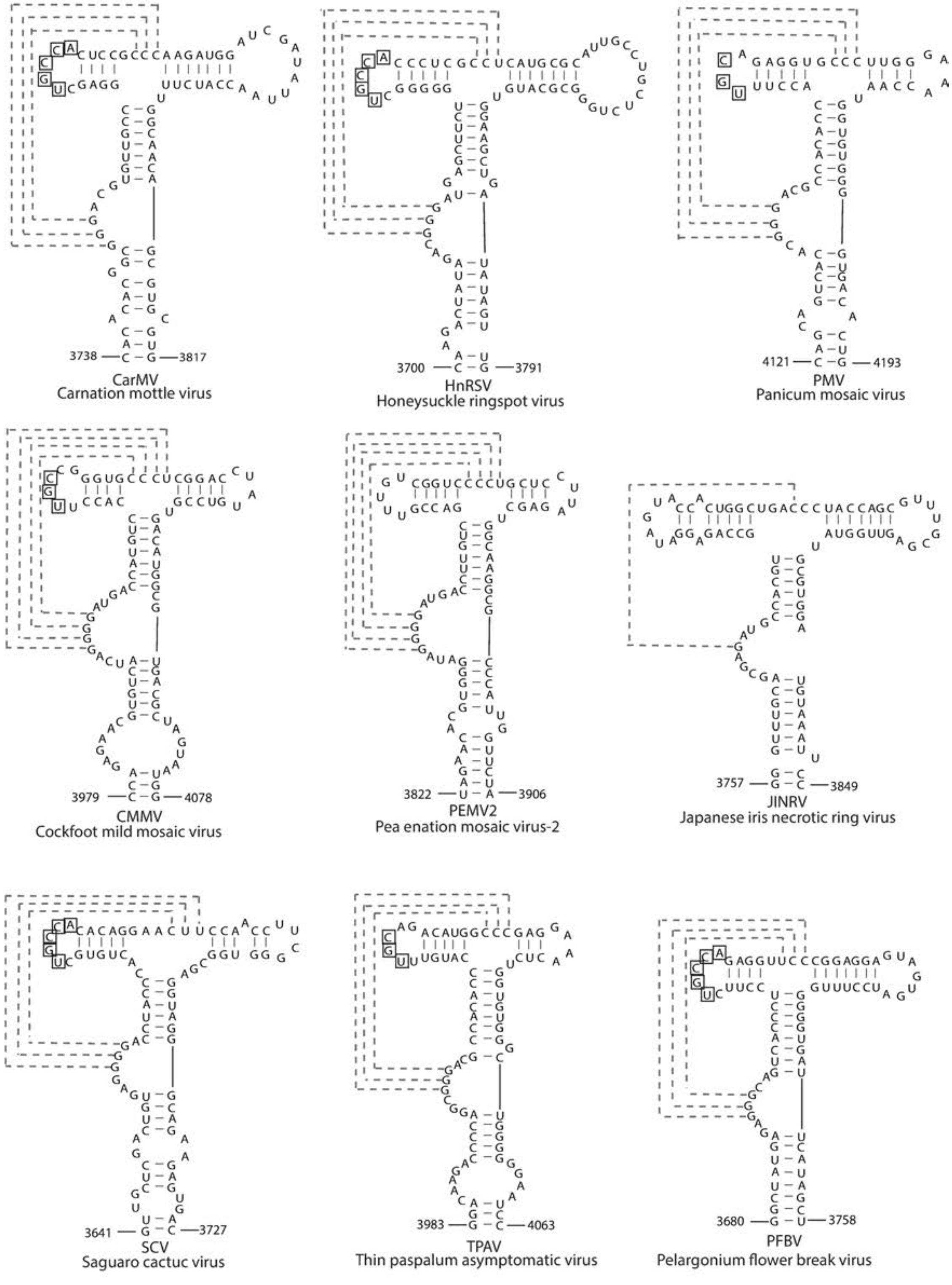


Supplementary Figure S2

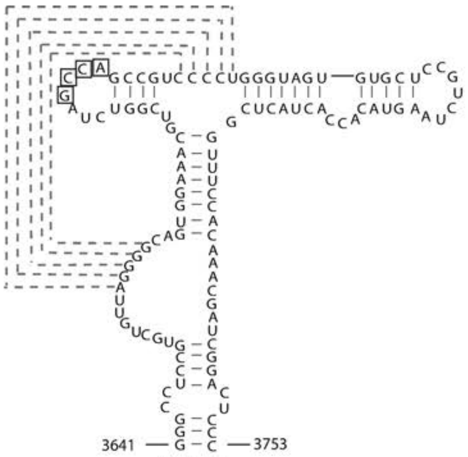


HCRSV
Hibiscus chlorotic ringspot virus

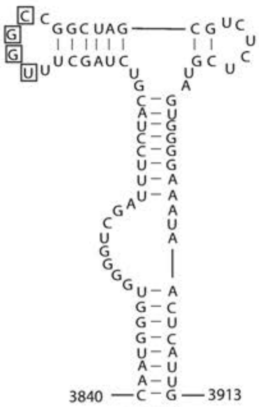


PSNV
Pea stem necrosis virus

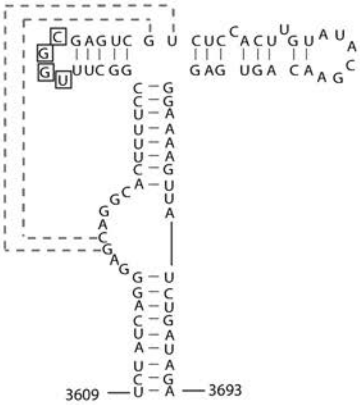


GaMV
Galinsoga mosaic virus

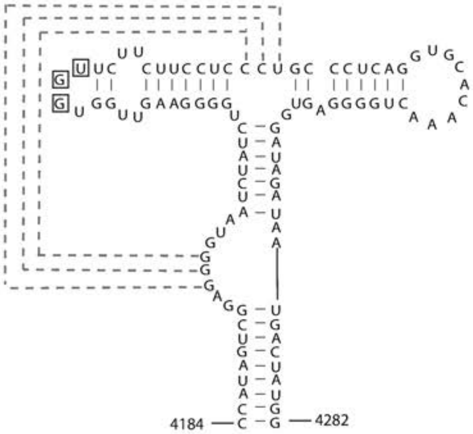


PoLV
Pothos latent virus

Supplementary Figure S2 (Continued)

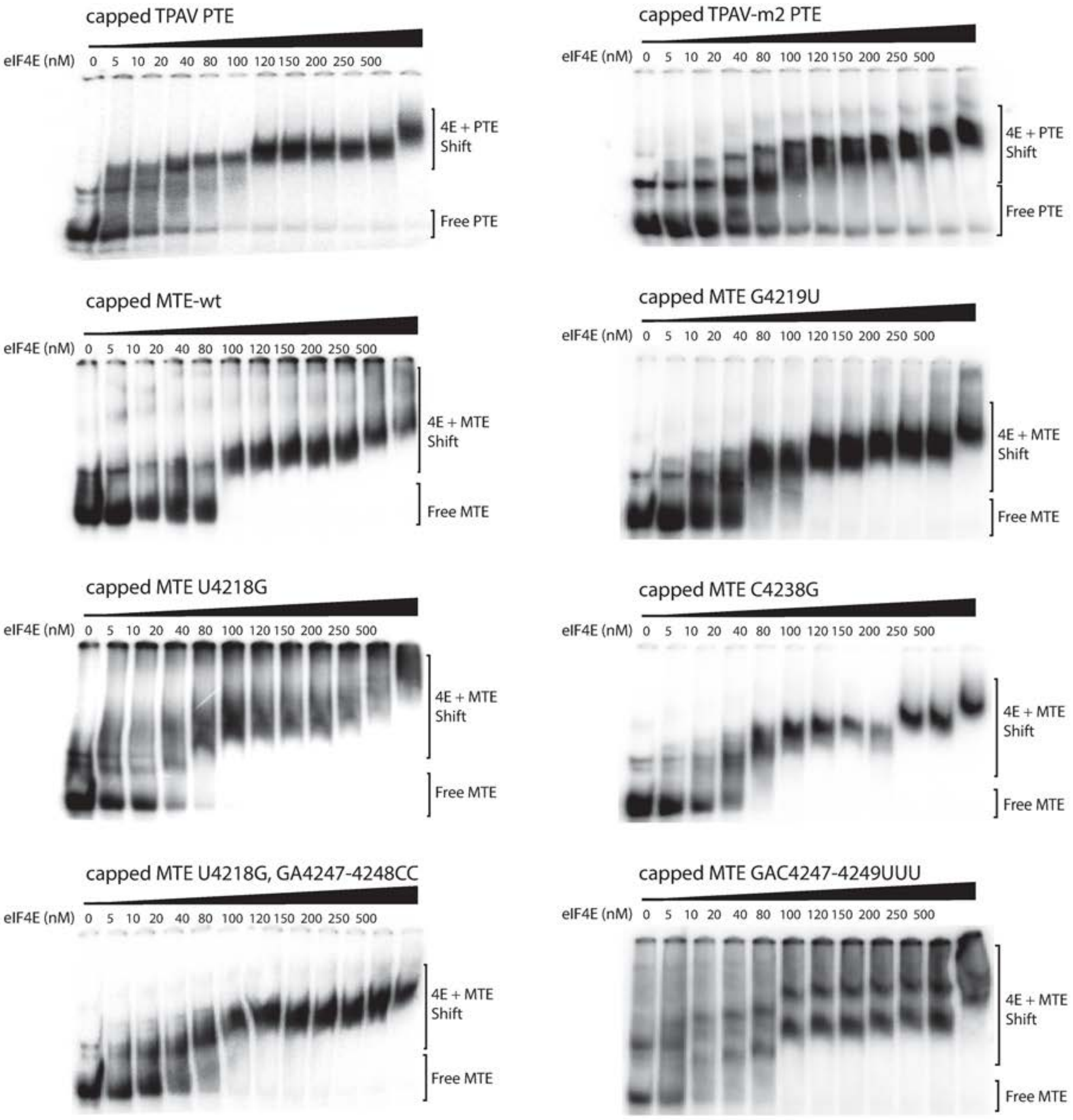


### Supplementary S3

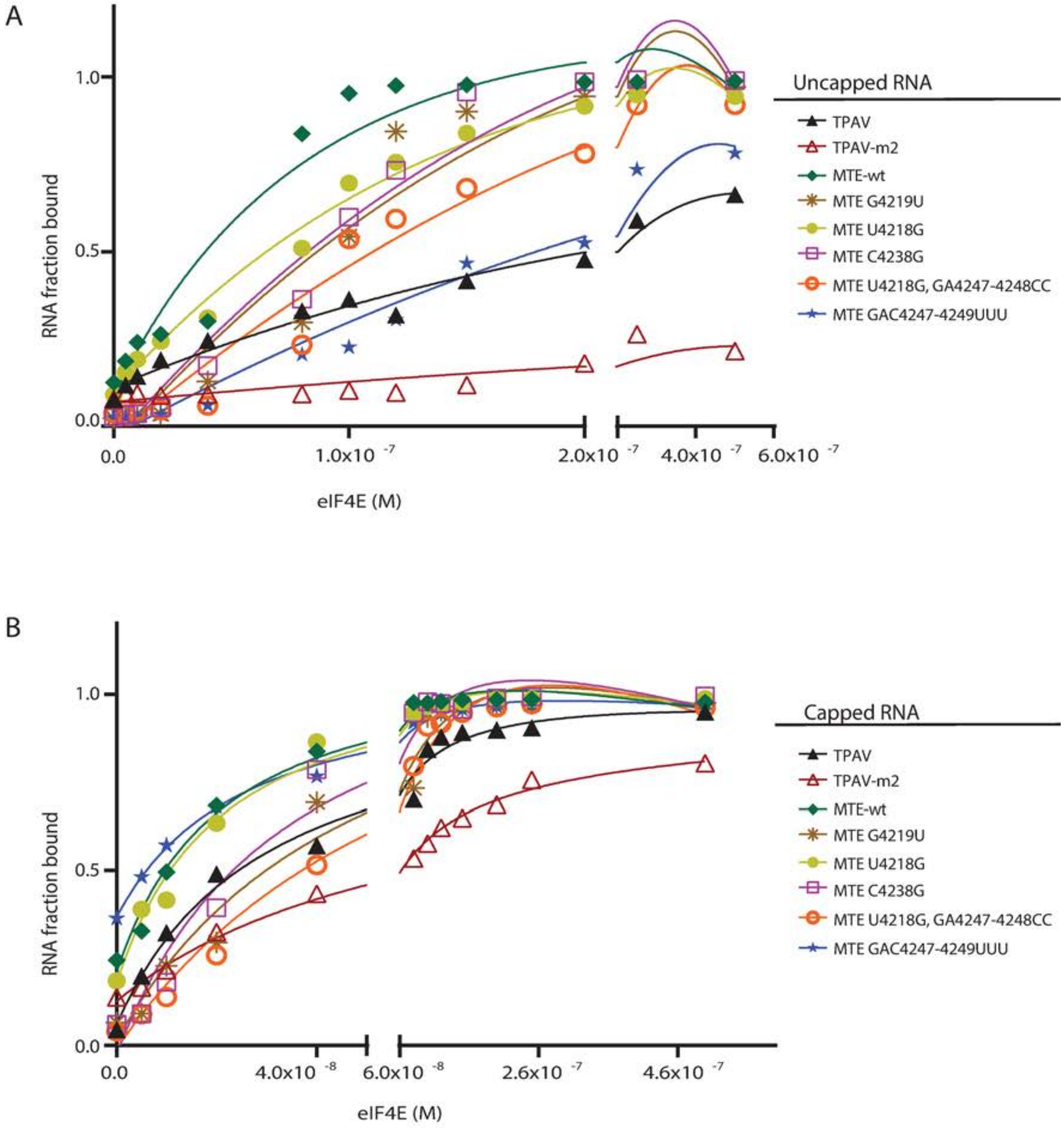


### Supplementary S4

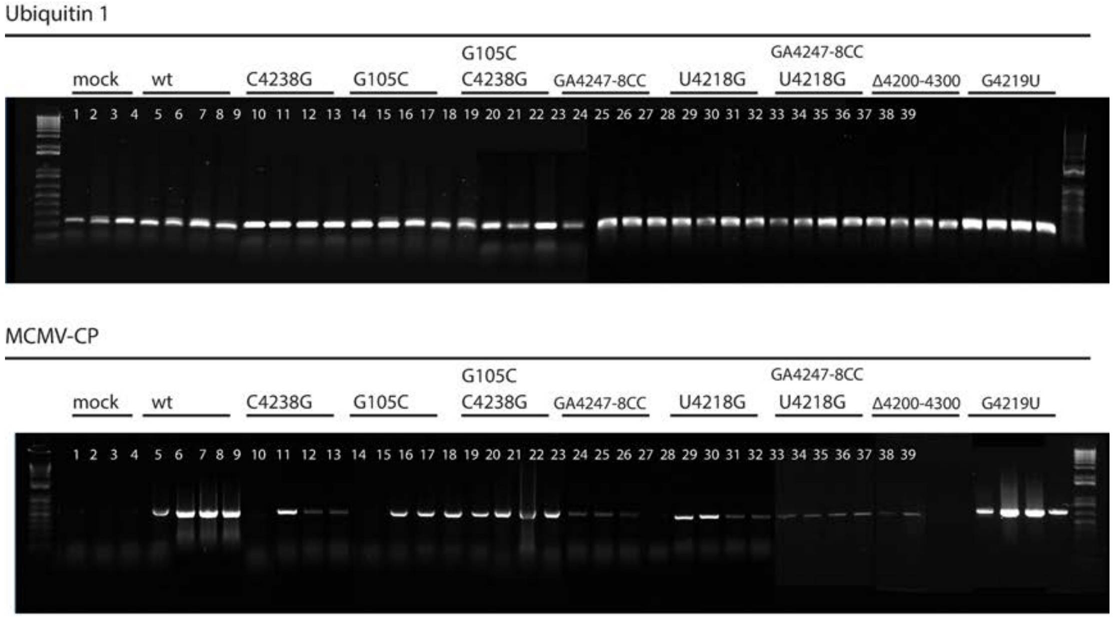


### Supplementary Figure S5

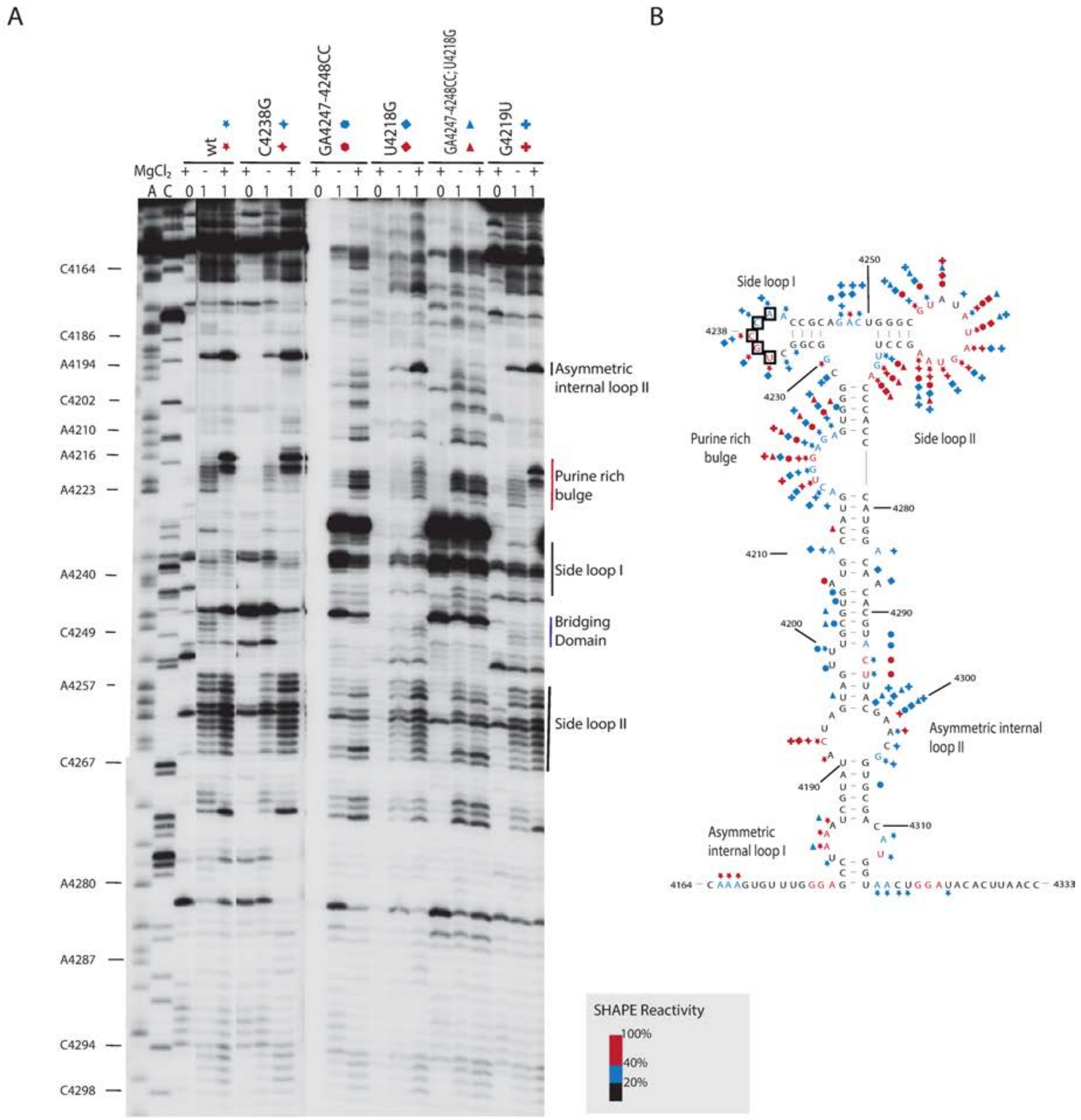


### Supplementary Figure S6

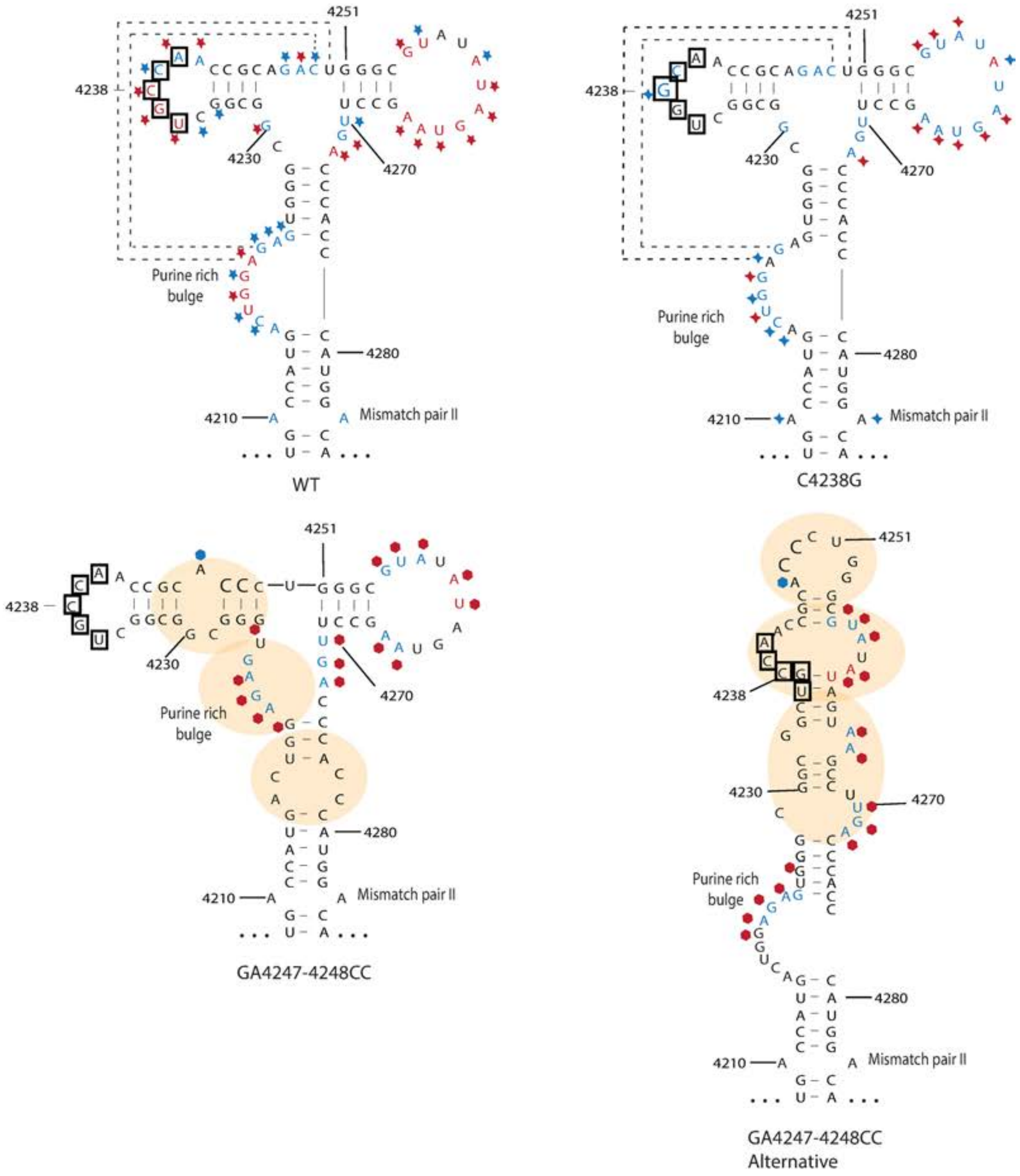


### Supplementary Figure S7

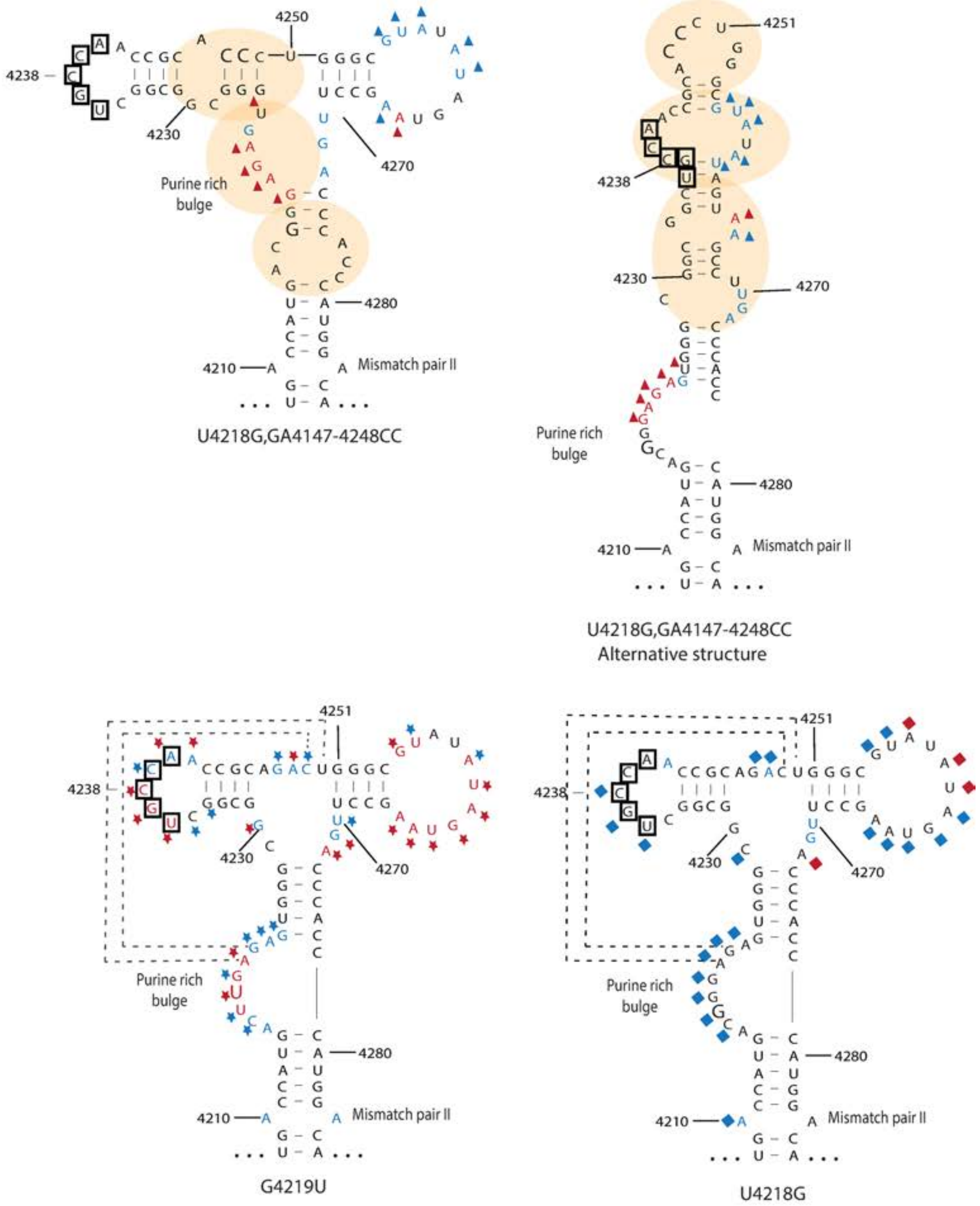


Supplementary Figure S7 (Continued)
